# Genomic analysis of the tryptome reveals molecular mechanisms of gland cell evolution

**DOI:** 10.1101/645952

**Authors:** Leslie S. Babonis, Joseph F. Ryan, Camille Enjolras, Mark Q. Martindale

## Abstract

Understanding the drivers of morphological diversity is a persistent challenge in evolutionary biology. Here, we investigate functional diversification of secretory cells in the sea anemone *Nematostella vectensis* to understand the mechanisms promoting cellular specialization across animals. We demonstrate regionalized expression of gland cell subtypes in the internal ectoderm of *N. vectensis* and show that adult gland cell identity is acquired very early in development. A phylogenetic survey of trypsins across animals suggests this gene family has undergone numerous expansions. We reveal unexpected diversity in trypsin protein structure and show that trypsin diversity arose through independent acquisitions of non-trypsin domains. Finally, we show that trypsin diversification in *N. vectensis* was effected through a combination of tandem duplication, exon shuffling, and retrotransposition. Together we reveal that numerous evolutionary mechanisms drove trypsin duplication and divergence during the morphological specialization of cell types and suggest the secretory cell phenotype is highly adaptable as a vehicle for novel secretory products.

## Introduction

The development of new tissue layers provides the opportunity to spatially segregate cell types enabling the compartmentalization of different functions. Cnidarians are diploblasts, comprised of an internal endodermal epithelium separated from an external ectodermal epithelium by a largely acellular matrix called mesoglea. Anthozoans (corals, sea anemones, and their kin) are unusual among cnidarians in their possession of internal tissues (pharynx and mesenteries) that arise by secondary epithelial fold morphogenesis following completion of gastrulation (1). Additional growth and differentiation of both internalized layers results in the morphogenesis of the pharynx and mesenteries and in an adult form quite different from that of medusozoans. In anthozoans, both layers (endoderm and ectoderm) are in contact with the gastric cavity, whereas in medusozoans (and, indeed, most other animals), the gastrovascular cavity is lined only by endoderm.

*Nematostella vectensis,* the starlet sea anemone, has become a valuable model for studies of animal body plan evolution (2–6); yet, little is known about the extent of cell diversity in the tissues that comprise the pharynx and mesenteries. The endodermal component of the mesenteries houses the germ cell precursors and two types of muscle cells and the few recent studies of the mesenteries in *Nematostella* have focused largely on these endodermal functions (7–9). The ectodermal component of the mesenteries is known to be populated by cnidocytes and gland cells (10) and two recent studies demonstrated the expression of multiple proteases in the mesenteries of *N. vectensis* (11, 12). Trypsins are the largest family of proteases, and although they have diverse functions, most trypsins are secreted to the extracellular environment and are therefore expressed in zymogen-type gland cells (13). A previous study cataloging trypsin diversity from prokaryotes and eukaryotes identified 75 trypsins in the genome of *N. vectensis* (14), suggesting that the few cell types identified anatomically as zymogen gland cells (10) may belie the digestive capacity of the mesenteries.

We sought to understand the evolutionary mechanisms promoting functional diversification at the cell and tissue levels in the mesenteries of *N. vectensis,* and to characterize the evolutionary history of a large (super)family of proteases expressed abundantly in the mesenteries. Building on a previous study using RNA-seq to characterize the expression profile of the mesenteries in *N. vectensis* (11), we show that the continuous epithelium comprising the internal ectoderm in *N. vectensis* is functionally partitioned into different regions associated with distinct morphologies and functions. Additionally, we show numerous lineage-specific expansions of trypsins and that trypsin diversification arises through novel domain acquisition. Finally, we propose a model by which the expansion of trypsins may have promoted specialization of gland cell subtype in cnidarians.

## Results

### Morphology and function of the internal ectoderm

We examined the fine structure of the internal and external ectoderm in the region of the mouth of *N. vectensis* during feeding for evidence of morphological and functional variation (Figure 1). Cells in the external ectoderm around the mouth are organized into a low cuboidal type epithelium that covers the closed mouth between feeding events (Figure 1A-D). In the presence of prey, the pharynx is partially everted, exposing the tall columnar epithelium of the pharyngeal ectoderm (Figure 1E-G). After passing through the pharynx (Figure 1H), ingested prey remains in contact with the ectodermal portion of the mesenteries, which is populated by cnidocytes and gland cells (Figure 1I-K).

**Figure 1.**
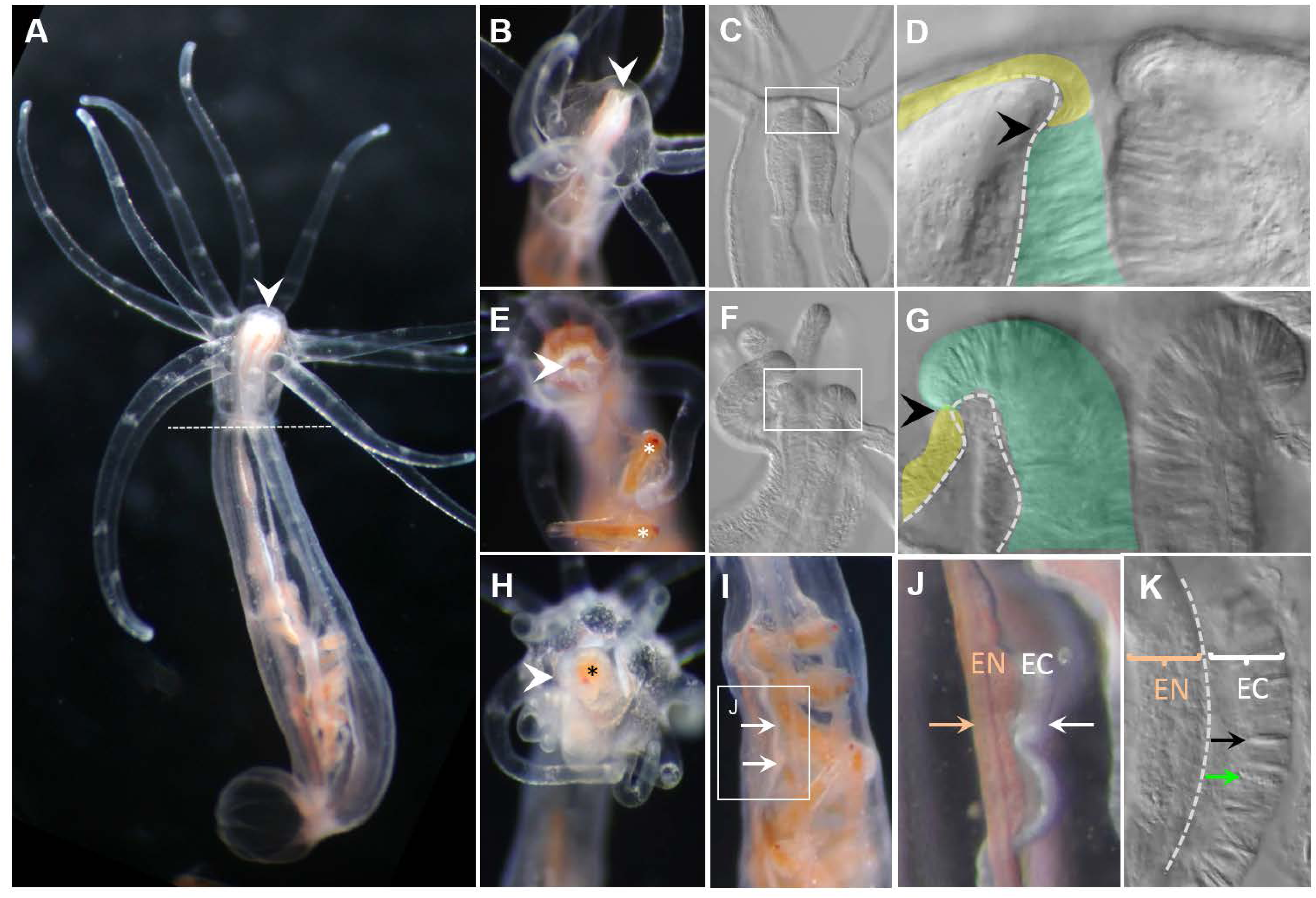
Morphology of the pharynx and mesenteries. (A) Adult polyp; the pharynx/mesentery transition is denoted by the dotted line. (B-D) Polyp at rest; the pharyngeal ectoderm (green) is retracted inside the oral ectoderm (yellow). (E-G) Partial eversion of pharynx occurs during capture/handling of prey (*Artemia* sp., indicated by *). (H-J) Ingested prey passes through the mouth and pharynx and remains in contact with the ectoderm (white arrows) of the mesenteries during digestion; colored arrow indicates endoderm of mesenteries (pigmentation from consumption of *Artemia*). (K) Cnidocytes (black arrow) and gland cells (green arrow) are restricted to the ectoderm of the mesenteries. C,D,F,G,K are DIC micrographs. D,G are false colored. E,H are oral views, the remaining images are lateral views. Dotted lines in D,G,K denote position of mesoglea. White arrowheads point to the mouth, black arrowheads denote transition from external to internal ectoderm.

The pharyngeal ectoderm is populated by numerous distinct electron dense (zymogen-secreting) and electron lucent (mucus-secreting) gland cells (Figure 2A-F). The adjacent non-secretory cells in this epithelium have distinctive apical electron-dense vesicles (Figure 2F). The proximal region of the mesentery (adjacent to the body wall) is comprised of endoderm while the distal portion (the free edge) is comprised of ectoderm (Figure 2G). The ectodermal region gives rise to both the cnidoglandular tract at the most distal extent (Figure 2H-J, M-O) and the ciliated tract more proximally (Figure 2K,L). Thin sections of the ectodermal mesentery in the oral region (near the pharynx) show abundant zymogen gland cells (Figure 2H-J), some of which contain secretory vesicles with heterogeneous contents (Figure 2J). Ciliated tracts are short and are present only in the oral end of each mesentery. Cells of the ciliated tract are highly proliferative and have apical motile cilia but do not have other distinguishing features (Figure 2K,L). The aboral mesentery lacks a ciliated tract but the cnidoglandular tract still contains numerous distinct zymogen gland cells, some with motile apical cilia (Figure 2M-O). Mucus-secreting cells were found in the pharyngeal ectoderm (Figure 2D) and in the external ectoderm of the body wall and tentacles (Supplemental file 1new), but never in the endoderm.

**Figure 2.**
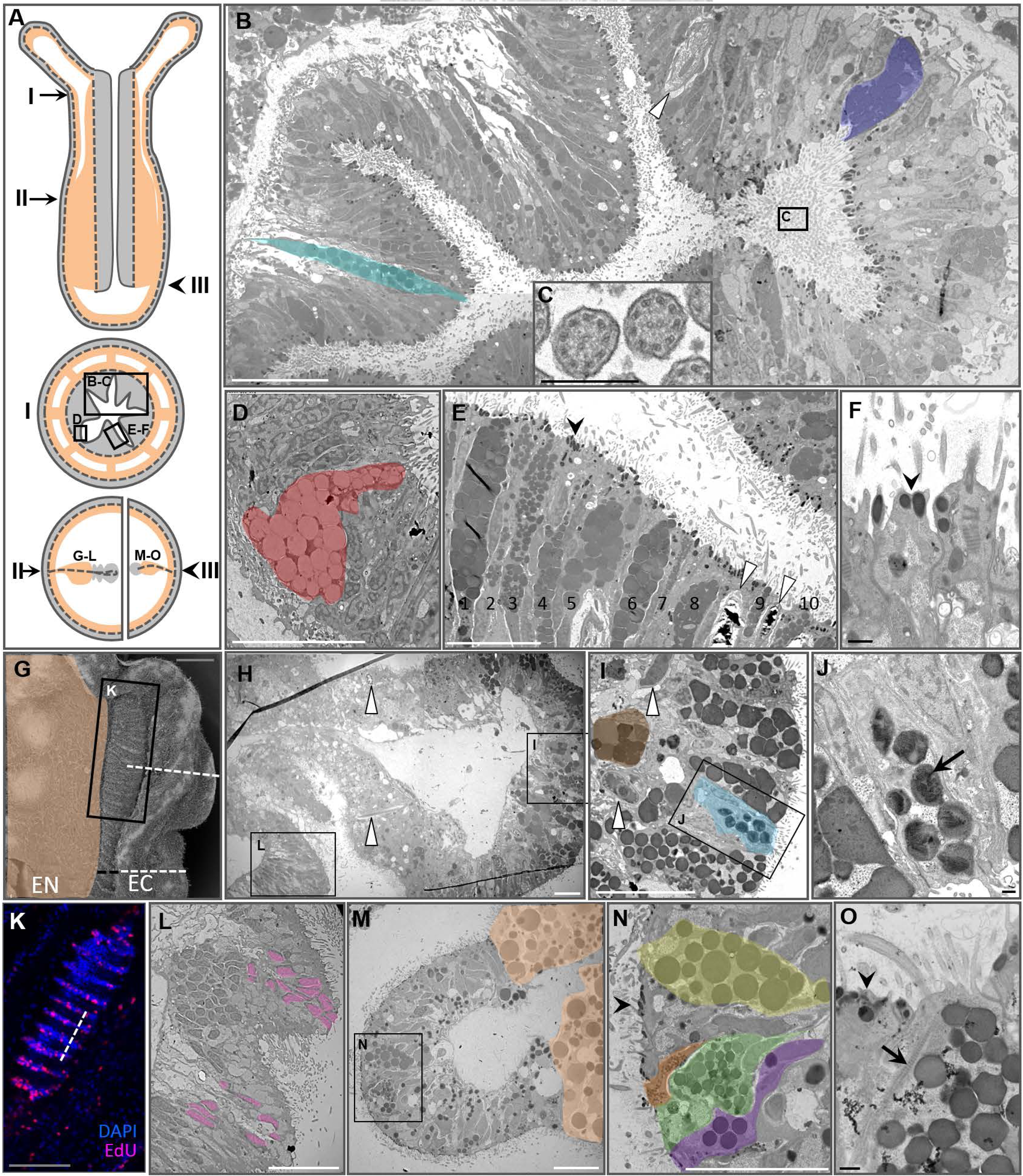
Fine structure of the pharyngeal ectoderm and ectodermal mesenteries. (A) Diagrammatic representation of regions where thin sections were examined through the pharynx (I) and at two positions in the mesenteries (II, III). Top – sagittal view of a primary polyp, middle – cross-section at position I, bottom – partial cross sections at positions II (left) and III (right). (B-F) TEMs of a region of the pharyngeal ectoderm (I) indicated by the box in A. (B) Two zymogen gland cells are false colored for emphasis. (C) Cross sections of cilia emerging from the pharyngeal ectoderm into the pharyngeal canal. (D) Mucus-secreting cell, false colored. (E) Ten zymogen-type gland cells and two cnidocytes. (F) Electron-dense vesicles (black arrowheads in E,F,N,O) in the apex of non-glandular ectodermal cells. (G) SEM of a mesentery from position II in panel A. The ectodermal (EC) portion has two parts: the ciliated tract (black line) and cnidoglandular tract (white line); the endodermal (EN) portion is false colored. (H-J) TEMs of the cnidoglandular tract of a mesentery from position II in A. Sections correspond to the position of the dotted line in G. (I) One zymogen type gland cell is false colored. (J) Some zymogen vesicles have heterogeneous contents (black arrow). (K) 3D reconstruction of a confocal z-stack through the ciliated tract of a mesentery from position II in A; pink – nuclei of proliferating cells (EdU), blue – quiescent nuclei (DAPI). (L) TEM section through the ciliated tract at the position indicated by the dotted line in K and by the box in H. Nuclei corresponding to panel K are false colored pink. (M-O) TEMs of a mesentery lacking a ciliated tract, position III in panel A. (M) Endoderm is false colored. (N) Zymogen gland cells are false colored. (O) Ciliary rootlet (black arrow) in the apex of a zymogen gland cell. Scale bars: white line – 10 um, black line – 500 nm, grey line (panels G, K) – 50 um. Panels B and H are composites of multiple micrographs. White arrowheads indicate cnidocytes throughout.

### Proteolytic enzymes are expressed in the developing mesenteries

We previously identified numerous genes encoding different classes of proteases to be upregulated in the adult mesentery of *N. vectensis* (11). Using *in situ* hybridization we examined the spatial and temporal expression of 10 proteases of various classes during early development of the pharynx/mesenteries to understand the ontogeny of digestive function and the onset of terminal gut cell differentiation. All genes examined were expressed in individual ectodermal cells of the mesenteries at the primary polyp stage, just after metamorphosis (Figure 3A,B); two protease genes (NVJ_82725 and NVJ_83864) were also expressed in the pharyngeal ectoderm of the primary polyp. There was surprisingly little variation in the onset of protease expression, although serine proteases (trypsins) consistently exhibited expression in the early planula stage before differentiation of the presumptive pharynx and mesenteries (Figure 3B). Double fluorescent *in situ* hybridization for two metalloprotease genes (NVJ_88668 and NVJ_2109) indicates both co-expression of these two enzymes in few cells at the aboral end of the pharynx and independent expression of the two genes in distinct cells of the ectodermal mesenteries in the late tentacle bud stage (Figure 3C).

**Figure 3.**
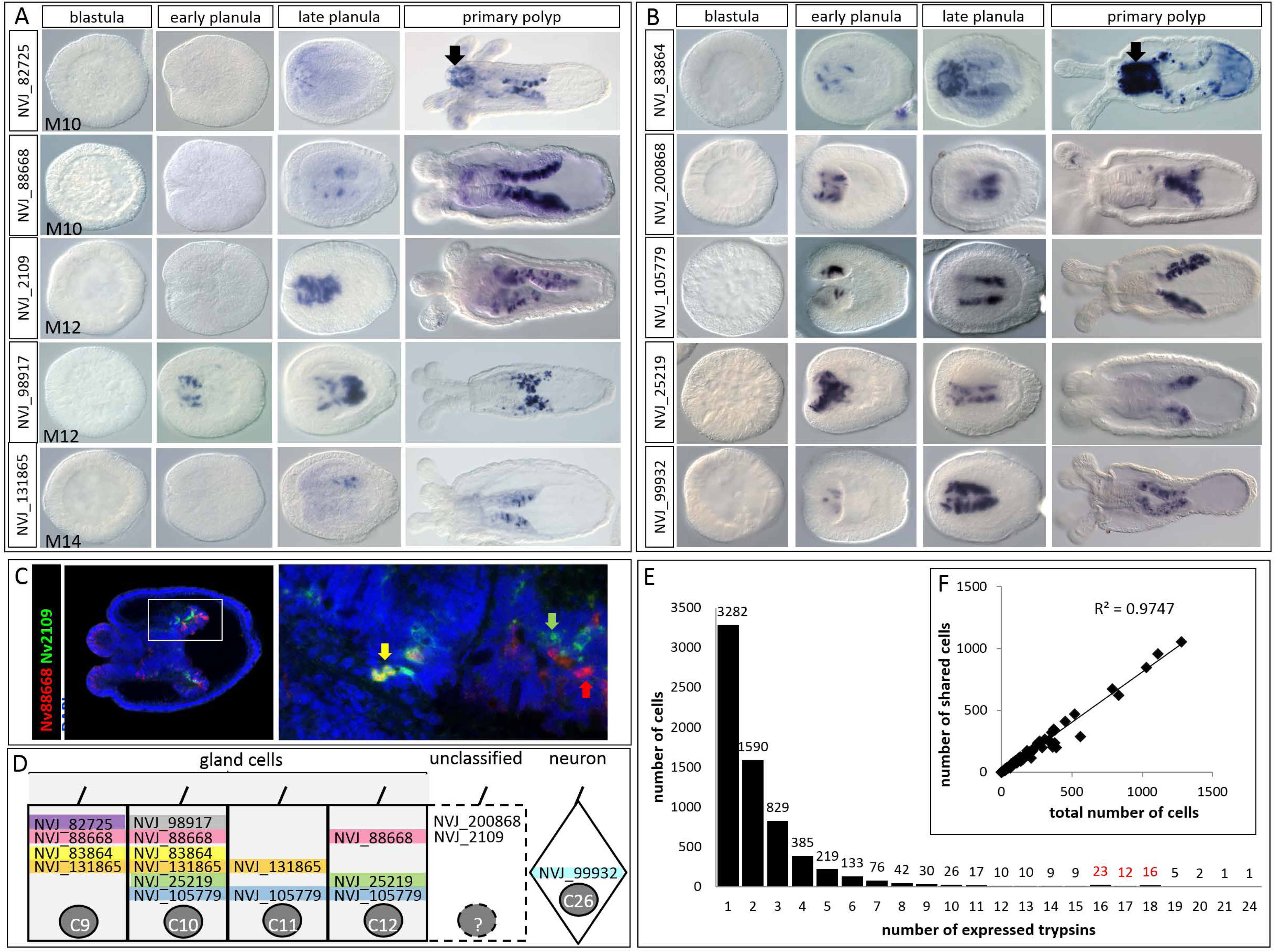
Expression of proteases in *N. vectensis*. (A-C) *In situ* hybridization of proteases in the (A) M family – metallopeptidases, astacins, and carboxypeptidases and (B) the S1 family – trypsins during early development. The oral pole is to the left in all images; black arrows indicate expression in the ectoderm of the pharynx. (C) Double fluorescent *in situ* hybridization showing the co-expression (yellow arrow) of two metalloprotease genes (Nv_88668 and Nv_2109) in the aboral ectoderm of the pharynx and independent expression of both genes in individual cells of the ectodermal mesentery (red and green arrows). Nuclei are labeled with DAPI. (D) Single-cell expression of genes from panels A and B in metacells identified by 1. Metacells C9-C12 are part of the “gland” cell cluster and C26 is part of the “neuron” cell cluster. Two genes (NVJ_200868 and NVJ_2109) are clearly expressed in the ectodermal mesenteries but were not reported by 1 (indicated by dotted lines). (E) Histogram of the number of expressed trypsins per cell (from 1); red font highlights a small second mode in the distribution. (F) Scatterplot of the number of cells exhibiting co-expression of multiple trypsins as a function of the number of total cells in which a particular trypsin is expressed.

The surprising lack of any obvious spatial segregation in protease expression led us to hypothesize that many proteases may be co-expressed together in the few anatomically distinguishable gland cells identified above (Figure 2). Using the raw data from a single-cell RNA-Seq study published previously (15), we show co-expression of six of the ten proteases we studied by *in situ* hybridization in a single putative gland cell (Figure 3D). Using the raw data from the same study and a very low cutoff for gene expression (N ≥ 1 read), we examined more fully the co-expression of the large superfamily of trypsin proteases and found 6,727 cells expressing at least one trypsin gene. Nearly 50% of the trypsin-expressing cells (3,282/6,727) appear to express only a single trypsin, while the remaining cells exhibited co-expression of up to 24 trypsins (Figure 3E). For each trypsin, we then examined the relationship between the ubiquity of expression (the total number of cells in which that trypsin is expressed) and the number of cells in which it is co-expressed with other trypsins and found a strong positive correlation (Figure 3F), confirming that the trypsins with the broadest expression profiles were most likely to be co-expressed with other trypsins.

### The tryptome is lineage-specific

To characterize the tryptome (all proteins with a trypsin domain) of *N. vectensis*, we searched the JGI gene models (https://genome.jgi.doe.gov/Nemve1/Nemve1.home.html) for all sequences containing a significant Trypsin or Trypsin_2 domain using hmmsearch (HMMER 3.1b2; http://hmmer.org) and constructed domain architecture diagrams for each protein (Figure 4). Of the 72 trypsin gene models that remained after curation (see Methods), 28 encode a trypsin domain but lack any other conserved domains and the other 44 encode a trypsin domain and at least one additional conserved domain. In total, trypsin domains were found in association with 24 other domains in *N. vectensis*. To determine if any of these associated domains were overrepresented in the tryptome, we compared the abundance of trypsin-associated domains in the tryptome and in the proteins predicted from the JGI gene models (N = 27,273 protein predictions). Six domains were found to be represented in high abundance (≥10%) in the tryptome: DIM, ShK, Lustrin_cystein, Sushi, MAM and SRCR (Figure 4A). The DIM and Lustrin_cysteine domains are present in low abundance throughout the predicted proteome (1 and 4 total domains, respectively), artificially inflating their perceived abundance in the tryptome. For ShK, Sushi, MAM, and SRCR, ≥15% of the domains found in the proteome were associated with trypsins.

**Figure 4.**
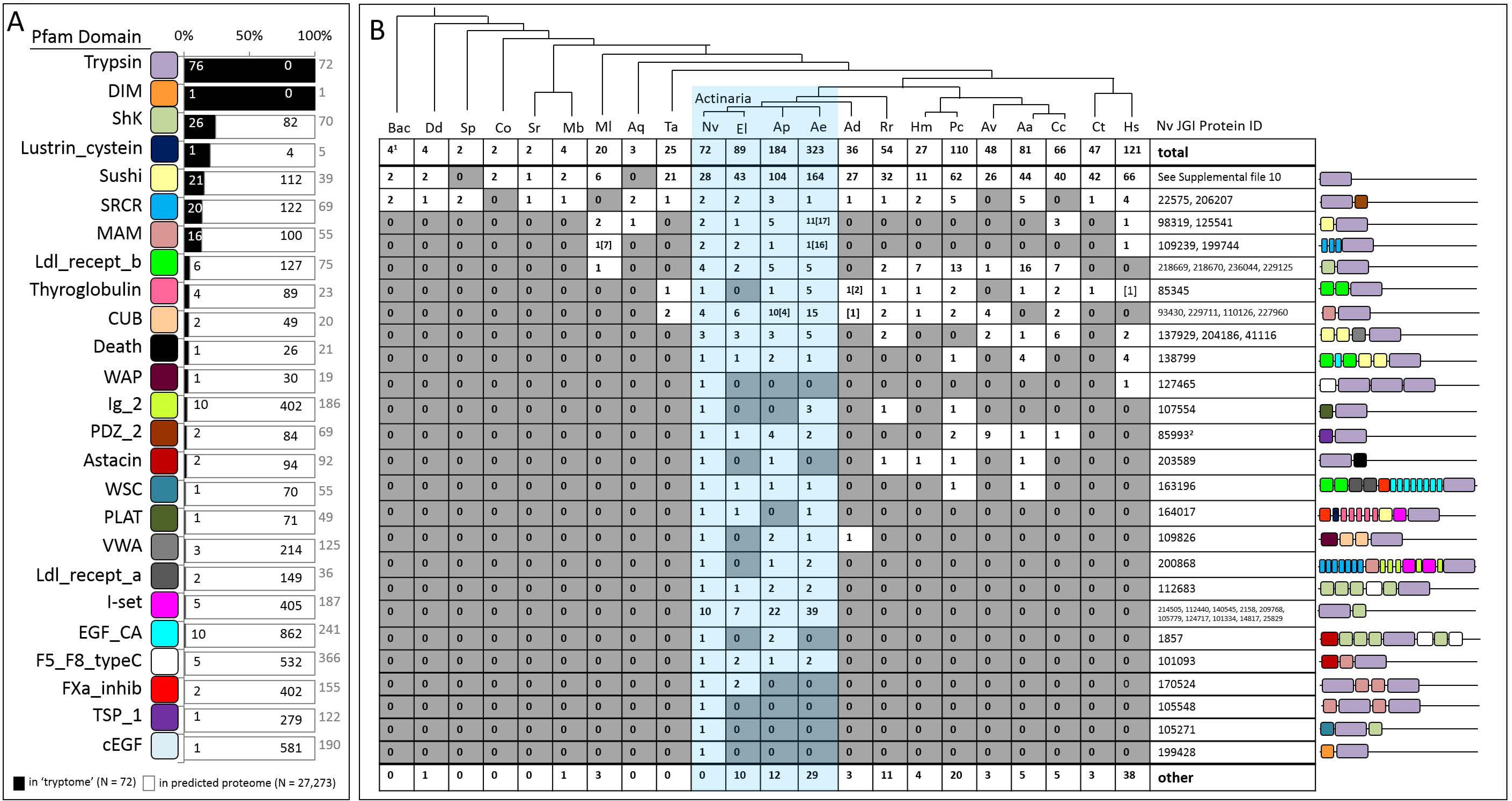
Characterization of the 72 proteins comprising the *N. vectensis* tryptome. (A) Abundance of the 24 trypsin-associated domains in the tryptome (black) and the remaining proteins in the predicted proteome (white). Numbers inside the bars reflect domain counts in both datasets; grey numbers to the right of the bars indicate total proteins containing the indicated domain. Pfam domains are listed on the left; see Supplemental file 4 for pfam domains IDs. (B) Phylogenetic distribution of trypsin domain architectures. Domain architecture models from the *N. vectensis* tryptome are shown on the right; proteins that differ only in the number of copies of associated domains were collapsed into a single row and these protein IDs are underlined. The number of proteins with identical domain composition are indicated for each taxon in the table. Actinarians (sea anemones) are highlighted in blue. ^1^Maximum number of trypsins from any one taxon in the bacteria/archaea database. ^2^El,Ae,Ap,Av have trypsin-TSP_1-ShK proteins which were included in this count. Bac – bacteria/archaea, Dd – *Dictyostelium discoidum*, Sp – *Schizosaccharomyces pombe*, Co – *Capsaspora owczarzaki*, Sr – *Salpingoeca rosetta*, Mb – *Monosiga brevicollis*, Ml – *Mnemiopsis leidyi*, Aq – *Amphimedon queenslandica*, Ta – *Trichoplax adhaerens*, El – *Edwardsiella lineata*, Ap – *Aiptasia pallida*, Ae – *Anthopleura elegantissima*, Ad – *Acropora digitifera*, Rr – *Renilla renilla*, Hm – *Hydra magnipapillata*, Pc – *Podocoryna carnea*, Av – *Atolla vanhoeffeni*, Aa – *Alatina alata*, Cc – *Calvadosia cruxmelitensis*, Ct – *Capitella teleta*, Hs – *Homo sapiens*

To determine whether the makeup of the tryptome was unique to *N. vectensis*, we searched for proteins with these same domain architectures in representatives from all domains of life (other cnidarians, bilaterians, non-metazoan eukaryotes, and a selection of prokaryotes). Two domain architectures were found to be present across taxa: those with only a trypsin domain, and those with a trypsin and a PDZ domain (Figure 4B). Trypsins appear to have expanded considerably after the origin of animals, as both choanoflagellate lineages had fewer than five trypsins but the ctenophore *Mnemiopsis leidyi* and the placozoan *Trichoplax adhaerens* (representing two of the earliest diverging animal lineages) both have at least twenty. Among the cnidarians, other actinarians (sea anemones) shared more trypsin proteins in common with *N. vectensis* than any other lineage; however, we identified three trypsins (NVJ_105548, NVJ_105271, and NVJ_199428) specific to *N. vectensis* that were absent even from *Edwardsiella lineata* (representing the genus sister to *Nematostella*).

### Trypsins diversified independently in cnidarians and bilaterians

To characterize the diversification of animal trypsins, we built a phylogeny of trypsin domains from taxa representing each of the five major animal lineages: bilaterians, cnidarians, placozoans, sponges, and ctenophores. Using this tree, we identify six clades of trypsins and classify them by their function in human: a non-catalytic group, the intracellular trypsins, tryptases and transmembrane trypsins, trypsins involved in coagulation and immune response, chymotrypsins, and the clade including granzymes, pancreatic trypsins, kallikreins, hepatocyte growth factors, and elastases (Figure 5A). Each of these includes representatives from bilaterians, cnidarians, and at least one placozoan, sponge, or ctenophore and likely represent the suite of trypsin clades present in the last common ancestor of animals. The *N. vectensis* tryptome includes representatives of 5 of 6 clades likely present in the common ancestor of animals. *N. vectensis* may have lost representatives of the tryptase/transmembrane clade as this these trypsins appear to be present in *M. leidyi*, *A. digitifera*, and bilaterians (Figure 5A). Proteins with only a trypsin domain were distributed throughout the tree, rather than being clustered in a single clade. Two proteins from *N. vectensis* (NVJ_203589 and NVJ_23745) had divergent trypsin domains and were detected only by the Trypsin_2 HMM, both of which appear to be part of the clade that includes trypsin-PDZ proteins (Supplemental file 2).

**Figure 5.**
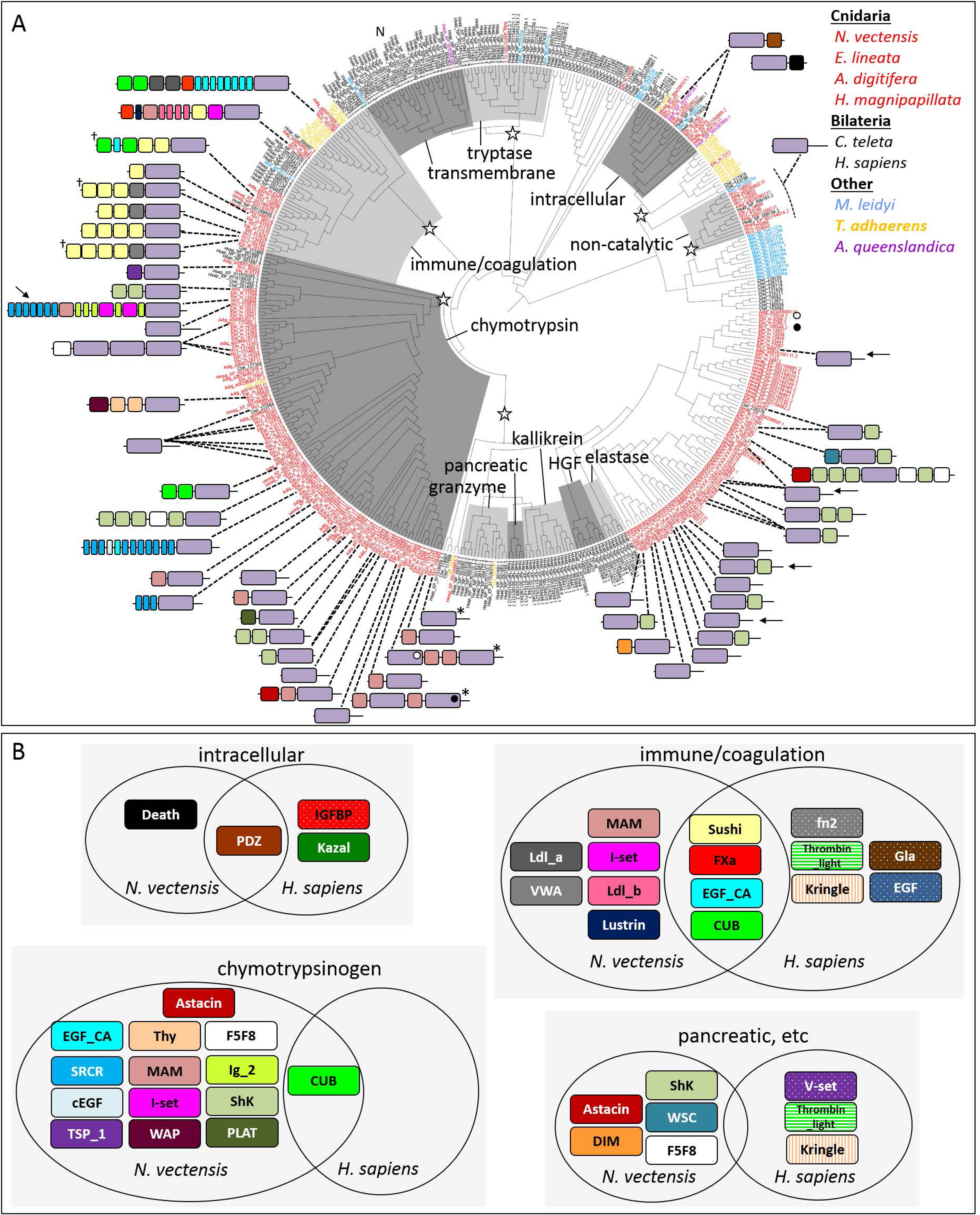
Evolutionary history of animal trypsins. (A) Phylogeny of animal trypsins; clades are named after the orthologs from *H. sapiens*. Domain architectures for *N. vectensis* proteins are shown. Proteins with multiple trypsin domains are polyphyletic; in such cases, the domain architecture diagram points to a single trypsin domain and the position of the other trypsin domains is indicated by open/closed circles. Open stars indicate clades that descended from single genes present in the last common ancestor of animals. N indicates the position of human neurotrypsin (NP_003610.2). The trypsin-death protein groups with trypsin-PDZ proteins but is not found on this tree. *N. vectensis* proteins characterized by *in situ* hybridization are indicated: arrows (this study), † (1), or * (1). (B) Comparison of the domain content of trypsins from *N. vectensis* and *H. sapiens*. The non-catalytic trypsins do not have associated domains and are not included. The tryptase/transmembrane trypsins are not represented in *N. vectensis* and are also not included. The “pancreatic” group includes granzyme, kallikrein, HGF, and elastase.

We compared the distribution of conserved domains from different clades of trypsins in *N. vectensis* and *H. sapiens* (Figure 5B). In *N. vectensi*s, domain diversity is greatest among the trypsins that group with human chymotrypsins, as trypsin domains from this clade co-occur with 14 associated domains. Chymotrypsins from *N. vectensis* share four domains in common with the immune/coagulation group, which is represented by 10 domains. The “pancreatic” group (including granzymes, kallikreins, HGF, and elastase) is characterized by only 5 domains, 3 of which are shared with chymotrypsins. Trypsins from the non-catalytic clade lacks associated domains and the intracellular clade uses unique domains (Death and PDZ). Four trypsin-associated domains (Sushi, EGF_CA, CUB, and FXa_inhibition) were found in the immune/coagulation clade of trypsins in both *N. vectensis* and *H. sapiens* and the PDZ domain is restricted to the intracellular clade of trypsins in both taxa. Surprisingly, there were no other domains found in common between *N. vectensis* and *H. sapiens* trypsins from the same clade. (See supplemental file 3 for distribution of human trypsin domain architectures.)

These analyses of trypsin distribution across animals revealed a surprisingly large tryptome in sea anemones (*N. vectensis* and *E. lineata*) relative to most other taxa (including another cnidarian, *Hydra magnipapillata*). To determine whether the tryptome of *N. vectensis* is reflective of other cnidarians, we built a phylogeny using representatives of each class within Cnidaria (Figure 6). We identify 16 clades of trypsins that include representatives of at least two lineages of anthozoans and two lineages of medusozoans, suggesting these clades may have been present in the stem cnidarian. Two clades (the trypsin-MAM and trypsin-ShK clades) seem to have undergone further expansion in anthozoans after their divergence from medusozoans.

**Figure 6.**
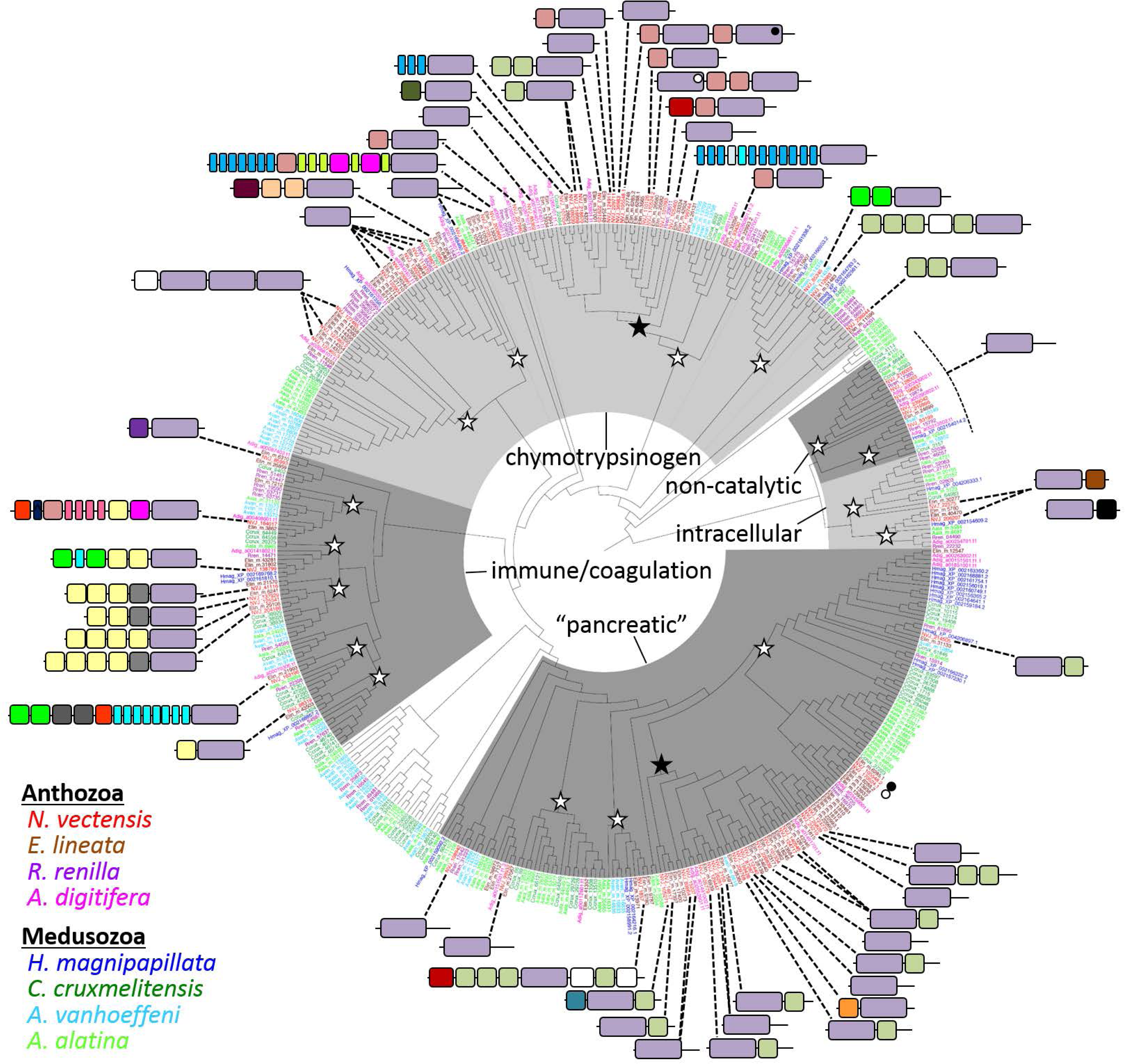
Gene phylogeny of cnidarian trypsin domains. Domain architectures for *N. vectensis* proteins are shown; domains are colored as shown in Figure 4A. Proteins with multiple trypsin domains are polyphyletic; in such cases, the domain architecture diagram points to a single trypsin domain and the position of the other trypsin domains is indicated by open/closed circles. Open stars indicate clades that descended from single genes present in the last common ancestor of cnidarians; closed stars represent possible anthozoan expansions. Clades are labeled as in Figure 5A; unshaded clades do not include any taxa from Figure 5 and cannot be resolved. The “pancreatic” group includes proteins that are sister to human granzyme, kallikrein, HGF, and elastase. Chymotrypsins are not monophyletic on this tree but are shaded together to facilitate comparison with Figure 5A.

### The Nematostella tryptome diversified through numerous mechanisms

To understand the mechanisms generating trypsin diversity in *N. vectensis*, we examined the evolutionary relationships of the 72 trypsin proteins in the tryptome (Figure 7A). Among the 72 predicted proteins, 85% (61/72) had all three conserved residues constituting the catalytic triad and are likely to function as proteases, 79% (57/72) were predicted to have a signal peptide and are presumably secreted, and 7% (5/72) were predicted to have a transmembrane domain (see Supplemental File 4). Four of the five clades of trypsins from *N. vectensis* (excluding the intracellular clade) include secreted trypsins, membrane-bound trypsins, and trypsins with divergent sequence that have likely lost their catalytic function. While most (70/72) of the trypsin domains were encoded across multiple exons (Supplemental file 5), two genes (NVJ_128003 and NVJ_216003) lack introns completely.

**Figure 7.**
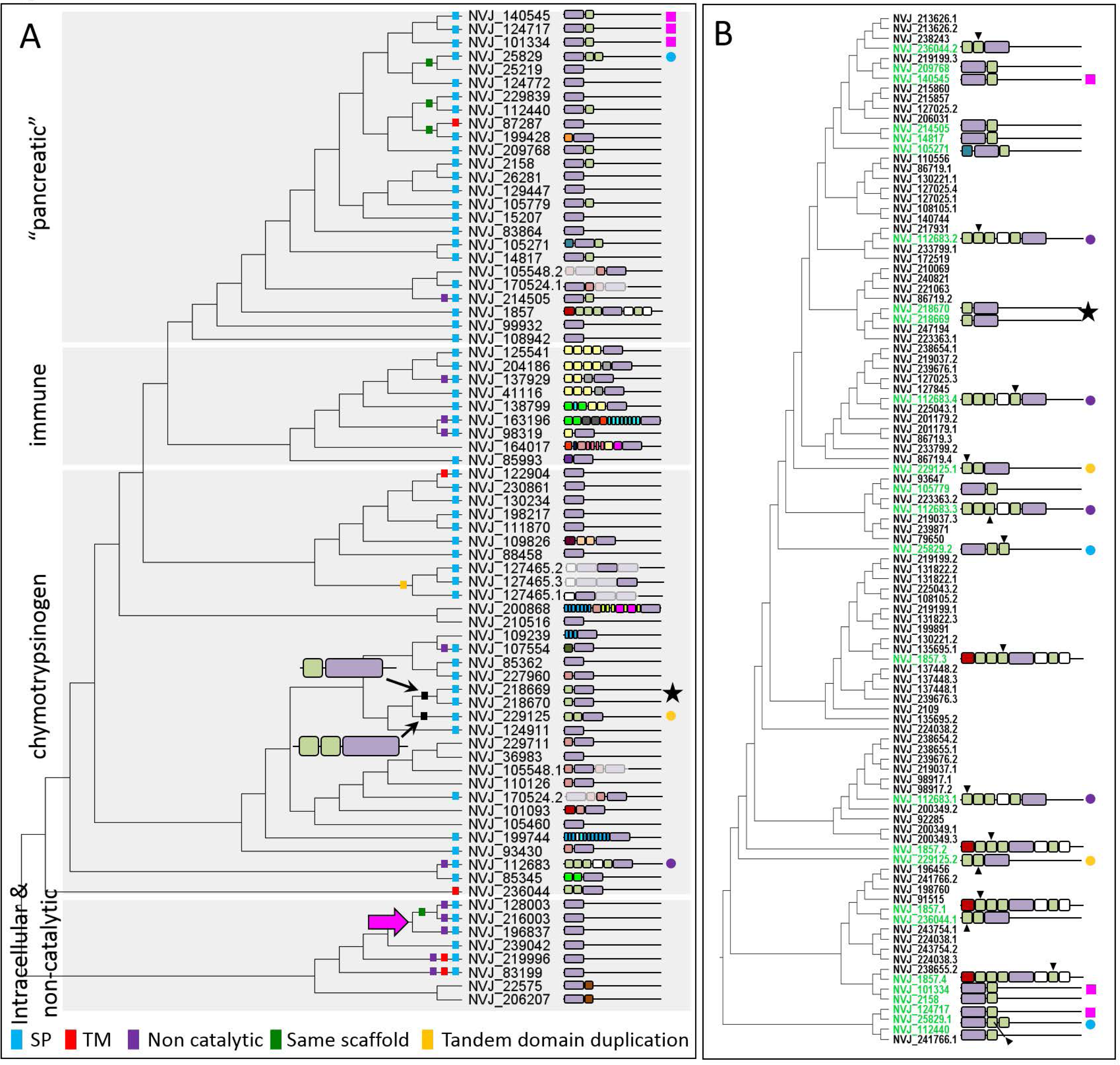
Relationships among trypsin and ShK proteins in *N. vectensis*. (A) Phylogeny of *N. vectensis* trypsin domains; domain architecture diagrams are shown to the right. Proteins with multiple trypsin domains appear multiple times in the tree; in these cases, the domain represented at each position in the tree is bolded. SP – signal peptide, TM - transmembrane domain. Trypsin domains lacking H-57, D-102, or S-195 are designated non-catalytic (see Supplemental file 4). Hypothesized ancestral states (models with dotted lines) are indicated in two positions for trypsin-ShK proteins; a revised hypothesis is indicated by the trypsin-ShK model with solid lines. (B) Phylogeny of *N. vectensis* ShK domains from the predicted proteome. Proteins with multiple ShK domains appear multiple times in the tree; in these cases, the domain represented at each position is indicated with an arrowhead. Proteins that have both trypsin and ShK domains are colored green. Colored symbols to the right of the domain architectures are provided to facilitate comparisons of identical proteins in panels A and B. Domains are colored as shown in Figure 4A.

Numerous trypsins from the “pancreatic” and chymotrypsin clades were associated with ShK domains. Likewise, over 30% (26/82) of the ShK domains in *N. vectensis* are associated with trypsins (Figure 4A). To determine if the combination of the trypsin and ShK domains may have duplicated together, we built a phylogeny of all 108 ShK domains from the *N. vectensis* proteins predicted from gene models (Figure 7B). Despite the abundance of trypsin-ShK associations, the distribution of trypsin-ShK proteins suggests that this domain combination may have duplicated together only once, giving rise to NVJ_218669 and NVJ_218670; Figure 7A).

### Trypsin diversity increases through new associations with old domains

Gene age can be estimated using a phylostratigraphic approach; in such analyses, the minimum age of a gene is inferred by identifying the last common ancestor in which the gene is present (16, 17). We examined the age of the trypsins found in *N. vectensis* and the age of each associated domain across all domains of life to understand the evolution of trypsin diversity. Trypsin-PDZ and a subset of the trypsin-only proteins likely arose before bacteria/archaea split from eukaryotes, over 2 billion years ago (Figure 8). While trypsin-only proteins are present in every lineage examined, trypsin-PDZ proteins appear to have been lost in several taxa including *C. owczarzaki*, *M. leidyi*, *A. vanhoeffeni*, and *C. cruxmelitensis* (Figure 4). All other associations between trypsin and other conserved domains appear to have originated after the stem metazoan diverged from the rest of life (∼800 million years ago). Many of the trypsin-associated domains originated long before they became associated with trypsin; for example, the astacin domain was present in the ancestor of all life but the trypsin-astacin association likely did not arise until the origin of Cnidaria (Figure 8A). By contrast, the SRCR domain and its association with trypsin likely arose in the stem metazoan as trypsin-SRCR proteins were found in *M. leidyi* (Supplemental file 6).

**Figure 8.**
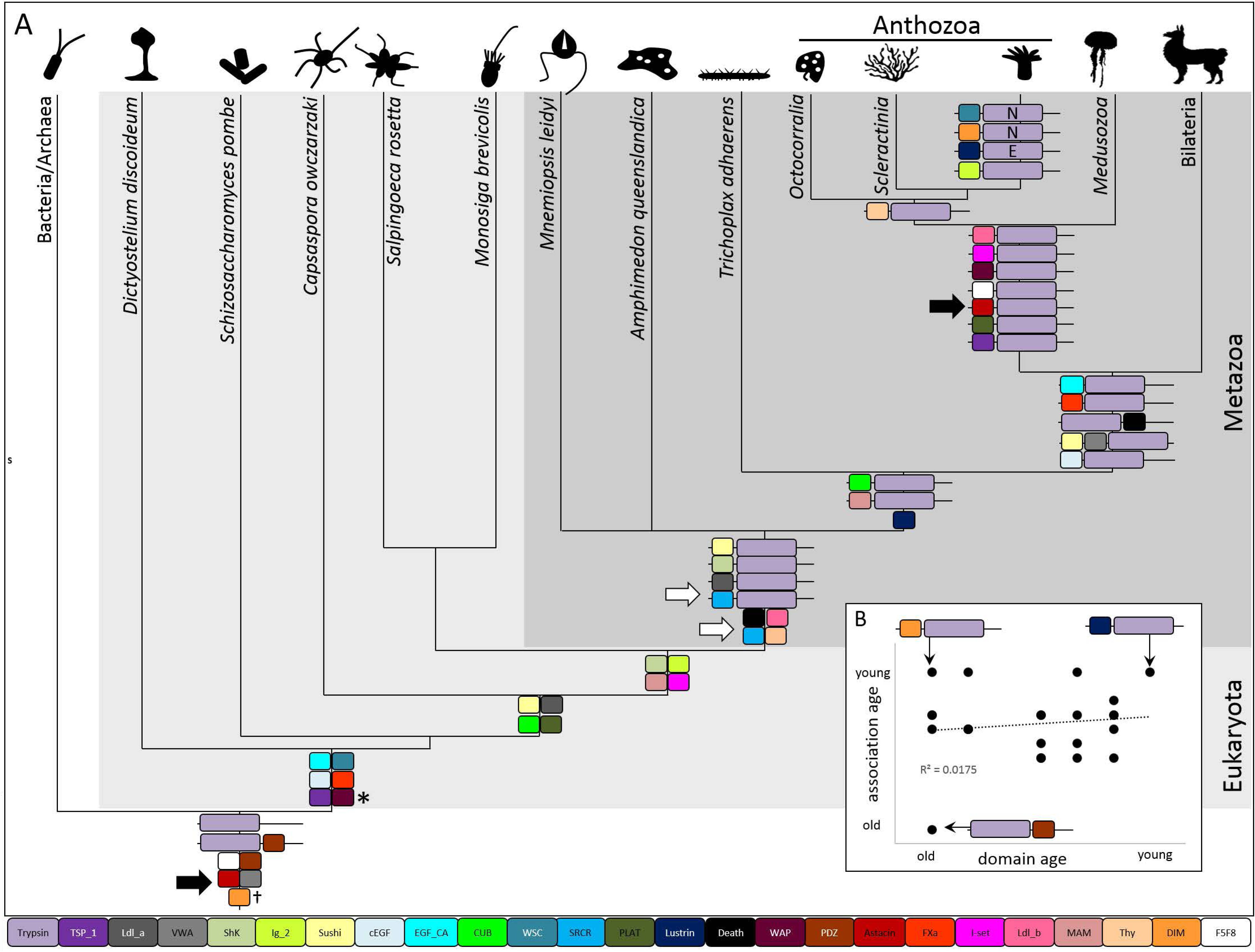
Evolutionary history of the tryptome. (A) Domain architectures at ancestral nodes indicate the origin of each domain combination; domains are reported at nodes where they appear for the first time with an independent (domain-specific) E-value ≤ 0.05. The origin of the SRCR domain and the origin of its association with trypsin are indicated by white arrows; black arrows indicate the origin of the astacin domain and the association between astacin and trypsin. E – Edwardsiidae only (*Edwardsiella* + *Nematostella*), N – *Nematostella* only. WAP domains (*) either evolved twice (in *D. discoideum* and the ancestor of animals) or this domain was lost in the intervening lineages. The DIM domain (†) has a complex distribution in bacteria, fungi, cnidarians, and insects. (B) There is no relationship between the phlyostratigraphic age of the domain and the age of its association with trypsin.

There does not appear to be a relationship between the age of the domain and the origin of its association with trypsin (Figure 8B). Two trypsin associations were found only in *N. vectensis*: trypsin-DIM (NVJ_199428) and trypsin-WSC (NVJ_105271), and one association was found only in *Edwarsiidae* (*Nematostella + Edwardsiella)*: trypsin-Lustrin_cystein (NVJ_164017). The WSC domain is present throughout eukaryotes (Figure 8A) but was associated with trypsin in only in *N. vectensis*. The Lustrin_cystein domain seems to have arisen in the last common ancestor of parahoxozoa (Placozoa + Cnidaria + Bilateria). These two associations represent extreme cases whereby trypsin diversity in *N. vectensis* arose through acquisition of both young (Lustrin_cystein) and old (WSC) domains.

## Discussion

### The not-so-simple cnidarian ectoderm

Although cnidarian body plans develop from only two tissue layers, morphological diversity varies widely across taxa. Similarly, only a dozen or so morphologically unique cell types have been described (10, 18), but cnidarian genomic and functional diversity rival that of any bilaterian lineage (4, 19). While the ectodermal layer comprising the external and pharyngeal epithelia may be contiguous, these regions are morphologically and functionally distinct in *N. vectensis* (Figure 1) (10, 12). In this study, we further demonstrate that the continuous layer of internal ectoderm from the pharynx through the mesenteries is equally heterogeneous. The pharyngeal ectoderm houses numerous zymogen and mucous cells while the ectoderm of the mesenteries houses only the former (Figure 2). This anatomical heterogeneity is supported by variable gene expression: some proteases are expressed throughout the pharyngeal and mesentery ectoderm, while others are restricted only to the mesentery ectoderm (Figure 3A,B) (also see (12, 20). Furthermore, the combinatorial expression of only two proteases can result in the development of at least three distinct cell types (Figure 3C). Together, the combination of a diverse tryptome and extensive trypsin co-expression (Figure 3E) suggests cell functional diversity in cnidarians may well exceed historical expectations.

We found no evidence of endodermal gland cells (zymogen type or mucous type) in our TEM or *in situ* hybridization results (Figure 2,3, Supplemental file 1). Indeed, all non-neuronal secretory cells (including mucous cells, zymogen cells, and cnidocytes), are restricted to the ectoderm in *N. vectensis* but their distribution is heterogeneous. Zymogen cell diversity, for example, is much higher in the internal than the external ectoderm (Figure 2, Supplemental file 1). This is consistent with the histological analyses of Frank and Bleakney (10) but seems to be in contrast with the distribution of gland cells in medusozoans. In *Hydra*, for example, zymogen gland cells are found exclusively in the endoderm (21). These observations suggest that the internalization of the ectoderm in anthozoans was a pivotal event in the diversification of specialized zymogen cells. Cell products secreted from the tentacle ectoderm may quickly become diluted in the water column, whereas the closed environment of the gastrovascular cavity limits the space over which secreted products can diffuse; thus, internalization created distinct selective pressures in different regions of the ectoderm. The selective pressure to secrete digestive enzymes into the enclosed gastrovascular cavity may have driven the development of gland cells in the internal ectoderm of anthozoans and the endoderm of medusozoans (and many bilaterians). As such, we see no reason to homologize the ectoderm of anthozoan mesenteries and the endodermal lining of the vertebrate midgut/pancreas (12). We consider it more likely that these tissues have converged on similar morphologies and gene expression profiles in response to similar selection pressures associated with extracellular digestion.

### N. vectensis trypsins have many putative functions

The trypsin domain catalyzes the cleavage of polypeptides at internal amino acid residues and is therefore essential for processing large proteins into smaller peptide chains. Digestive trypsins are synthesized in secretory cells with zymogen type secretory granules where they are packaged into vesicles for release into the gut. We show that there are at least 10 morphologically distinct zymogen gland cell types in the pharyngeal and mesentery ectoderm of *N. vectensis* (Figure 2). Further, we demonstrate that numerous proteases are expressed in the same tissues (Figure 3) and that the vast majority of trypsins in *N. vectensis* encode a signal peptide (Figure 7A). Using published single-cell expression data (15), we identified ten putative gland cells that express trypsins, at least two of which also express synaptotagmin (Supplemental file 4), which facilitates fusion of the vesicle with the cell membrane during regulated secretion. These combined features point to the expression of trypsins in secretory cells of the internal ectoderm and strongly support a role for trypsins in extracellular protein degradation in *N. vectensis*.

Numerous trypsins were expressed outside of the putative gland cells identified by Sebe-Pedros et al (15). At least 20 cells categorized by these authors as neural cells exhibited trypsin expression but unlike gland cells, the maximum number of trypsins expressed by any putative neuron is three (Supplemental file 4). We show trypsin-expressing cells differentiating very early in development, in the invaginating pharynx/mesenteries (Figure 3). Neurons expressing RFamide and Elav are also undergoing terminal differentiation in this tissue at this developmental stage (18, 22). Indeed, the trypsin protease NVJ_99932 (Figure 3) is co-expressed with two other trypsins (NVJ_230861 and NVJ_130234) in a putative neuron expressing GABA and dopamine receptors (Supplementary file 7). In vertebrates, secretion of neurotrypsin from the pre-synaptic membrane facilitates degradation of the extracellular matrix during synaptic plasticity and axon guidance (23). Although 17 different trypsins were expressed in putative neurons, none of the trypsins from *N. vectensis* clustered with human neurotrypsin (Figure 5), suggesting this function may have been acquired independently from different ancestral trypsins.

Trypsins are important regulators of tissue remodeling; as such, upregulation of trypsins and other proteases occurs coincident with wound healing and tissue regeneration (24). Recent studies of regeneration in *N. vectensis* demonstrated that a new pharynx will regenerate from the oral ends of the mesenteries after amputation (25) and that many proteases are expressed abundantly during this process (26). Thus, the mesenteries appear to play an important role in directing the tissue remodeling process in *N. vectensis*. In support of this, a study of wound healing in response to a body wall injury demonstrated that the mesenteries come into direct contact with damaged tissue during the healing process (27). This study also showed that two trypsins (NVJ_107554 and NVJ_112683) are among the top genes undergoing upregulation during wound healing in *N. vectensis*. While NVJ_112683 was not reported in the single-cell dataset, NVJ_107554 is expressed in two putative gland cells (metacells C12 and C19, Supplemental file 4). One of these (metacell C12) is also the site of expression of three proteases examined by *in situ* hybridization in this study (Figure 3). Thus, mesentery-expressed trypsins play important roles in the cell and tissue biology of *N. vectensis* during wound healing and regeneration and these roles may vary through ontogeny.

Beyond their roles in digestion and tissue remodeling, trypsins are an important component of the innate immune system. In vertebrates, immune trypsins play a role in blood coagulation and are part of the complement system which recognizes foreign particles (28). In symbiotic cnidarians, immune trypsins play a role in the beneficial interaction between the host and the alga (29). While *N. vectensis* does not host symbiotic algae, a previous study aimed an understanding the origin of the innate immune system reported the expression of three immune system trypsins in *N. vectensis*: MASP (NVJ_138799) and two paralogs of Factor B (NVJ_41116, NVJ_204186), each of which were expressed in the endoderm (gastrodermis) of juvenile polyps (30). We found that the two factor B orthologs were also co-expressed in a single putative gastrodermal cell (Supplemental file 4) further supporting a role for the endoderm in the immune response of *N. vectensis*. One trypsin (NVJ_127465) was not reported in the single-cell dataset (15) but was among the genes found to be significantly upregulated in the tissue-specific transcriptome of nematosomes, which may also play a role in the immune system of *N. vectensis* (11). This gene clustered with human chymotrypsin genes, not the immune system trypsins (Figure 5); as such it may have acquired a role in the immune system secondarily.

### Trypsin functional diversity has undergone numerous expansions

Two groups of trypsins (intracellular and non-catalytic) are found in all domains of life (Figure 4), both of which form monophyletic groups in animals (Figure 5). Trypsin diversity expanded rapidly with the origin of animals, as representatives of two of the earliest diverging animal lineages (ctenophores and placozoans) have at least 20 trypsins (Figure 4). The phylogeny of animal trypsins suggests the last common ancestor of animals may have had at least six major groups of trypsins (Figure 5). Sponges are unusual among animals in that they have only three trypsins – two trypsin-PDZ paralogs and a trypsin-Sushi protein (Supplemental file 6). This suggests either extensive loss of trypsins in Porifera or independent diversification of trypsins in ctenophores and in the stem of parahoxozoa. The evolutionary history of trypsin-Sushi, trypsin-SRCR, and trypsin-ShK proteins sheds little light on this topic; while all three associations are found in ctenophores, trypsin-Sushi proteins are missing from placozoans and trypsin-SRCR and trypsin-ShK proteins are missing from both sponges and placozoans (Figure 4,8, Supplemental file 6). Presently, it is difficult to determine if these associations were either lost in sponges and placozoans or arose independently in ctenophores and the stem of planulozoa. Considering the association between the ShK and trypsin domains also seems to have been lost in the bilaterian lineage (Figure 4, Supplemental file 6), we think independent gain of ShK domains in ctenophores and cnidarians is likely.

The ancestral cnidarian may have had a far more diverse suite of trypsins than the ancestral animal. Indeed, our data suggest there were at least 17 lineages of trypsins present in the last common cnidarian ancestor (Figure 6) and 12 of the associations between trypsin and another conserved domain in *N. vectensis* are specific to cnidarian lineages (Figure 8). Furthermore, while the diversity of trypsins in *N. vectensis* rivals that of *H. sapiens* (Figure 4), there is little conservation in the associated domains in these taxa (Figure 5B). Furthermore, within each clade of trypsins, sequences from cnidarians and bilaterians form distinct groups, suggesting that secretory cell function expanded independently in the cnidarian and bilaterian stem lineages. There was extensive divergence in the trypsin gene superfamily during the diversification of cnidarians but anthozoans seem to have undergone additional radiations in at least two trypsin clades. Anthozoans are the most speciose group of cnidarians and are largely sessile; thus, selection for trophic specialization and sympatric niche diversification may be stronger among anthozoans than medusozoans.

### Numerous evolutionary mechanisms contributed to the rise of trypsin diversity

Multidomain proteins are more common than proteins with only a single domain as domain recombination increases versatility in protein function (31). Selection to maintain the catalytic activity of the trypsin domain while allowing the context in which this domain is expressed to vary was a primary driver of diversification of this gene family. Surprisingly, there was little conservation in trypsin-associated domains across animals, even among cnidarians (Figure 4, Supplemental file 6), suggesting the trypsin domain underwent significant duplication before the diversification of cnidarians and that the associated domains have been continuously gained and lost in each cnidarian lineage. Furthermore, nearly 40% (28/72) of the proteins comprising the *N. vectensis* tryptome have only a trypsin domain (Figure 7) yet these trypsin-only proteins did not form a monophyletic group (Figure 5,6). These results suggest that trypsin domains themselves may be rapidly gained and lost from evolutionarily unrelated proteins, further underscoring the selective advantage of a trypsin domain.

Trypsins were associated with 24 other conserved protein domains in the *N. vectensis* tryptome, only few of which were over-represented in the tryptome (Figure 4A). The ShK domain is a short peptide found in a K-channel inhibitor originally isolated from the sea anemone *Stichodactyla helianthus* (32). The ShK phylogeny (Figure 7B) further suggests this domain is gained and lost easily, as ShK domains from sister trypsins were almost never monophyletic. Consistent with this, every ShK domain in the tryptome of *N. vectensis* was encoded by only a single exon (Supplemental file 5), supporting the possibility of rapid evolution through exon shuffling. Two trypsin-ShK proteins (NVJ_218669 and NVJ_218670) were found to be sister in both phylogenies, suggesting they arose by duplication of the combined domains. These two genes are encoded on the same scaffold and are separated by approximately 1000bp of genomic DNA; thus, they are likely the result of a recent tandem duplication event. What role the ShK domain plays when it is paired with the trypsin domain is not known but the overabundance of these two combined domains in cnidarian tryptomes (Supplemental file 6) combined with the multiple independent origins of this domain combination in *N. vectensis* (Figure 7B) suggest the pairing provides a strong selective advantage in the biology of cnidarians.

We also found evidence of trypsin diversification independent of the acquisition of associated domains. One gene (NVJ_127465) encodes three trypsin domains, all of which form a monophyletic group suggesting this gene structure arose through tandem duplication of the trypsin domain. Likewise, we identify four cases where sister trypsins are found on the same scaffold, suggesting tandem gene duplication. Additionally, two trypsins were found to lack introns (NVJ_128003 and NVJ_216003), suggesting these two arose through recent retrotransposition. These two genes are also on the same scaffold, suggesting retrotransposition may have been followed by tandem gene duplication. The tryptome from *H. sapiens* also includes two proteins with three trypsin domains each (Supplemental file 6). All six of these trypsin domains from *H. sapiens* are found in the tryptase/transmembrane clade (Supplemental file 3), whereas the three domains in NVJ_127465 group with chymotrypsins. Despite their similar domain architecture, triple-trypsin domain proteins appear to have evolved multiple times.

Diversification of the trypsin superfamily through gene duplication and divergence has been continuous, suggesting an important role for trypsins in the evolutionary success of animal lineages. Early expansions, before the diversification of animals may have promoted the origins of the innate immune system and extracellular digestion, facilitating the evolution of large body size and increased longevity in this lineage. Lineage-specific expansions (between genera, for example) in the digestive trypsins enable the development of specialized of taxon-specific tryptomes, which can support niche specialization in otherwise similar taxa (Figure 4,6,8). The expansion of the tryptome of *N. vectensis* was facilitated by gene duplication followed by the acquisition of additional domains (Figure 8). In some cases, the acquisition of an associated domain by a trypsin protein occurred soon after the origin of the acquired domain (e.g., trypsin-SRCR), while in other cases this acquisition occurred long after the associated domain first appeared (e.g., trypsin-astacin). Indeed, we found no relationship between the age of the domain and the age of the association with trypsin (Figure 8B), further supporting the idea that trypsin domain architectures diversify continuously and are not dependent on the origin of novel domains.

New genes are thought to arise rapidly *via* duplication and divergence (33) but the proportion of novel genes that become functionally integrated into signaling networks may be very small. The factors that promote retention of new genes following duplication are not well understood but may involve complimentary degenerative mutations promoting subfunctionalization in the duplicates (34). Under this framework, sister trypsins that result from a recent duplication event should be expressed in differing contexts, for example in different developmental stages or in different cell types. Consistent with this, analysis of the single-cell expression of NVJ_218669 and NVJ_218670, which appear to have undergone recent duplication, suggests that NVJ_218669 is expressed in a subset of the cells that express NVJ_218670 (Supplemental file 4). Only 51 of the 72 trypsins in *N. vectensis* were reported in the single-cell study (15), making a detailed analysis of shared expression patterns in sister trypsins premature; however, a targeted study of this protein superfamily in the future may reveal novel cis-regulatory regions, further enlightening the processes involved in the diversification of this protein family.

### Secretory cells and the evolution of cnidarian body plans

Resolving the embryological origin of cnidarian gland cells will be important for understanding the evolution of life history in Cnidaria. If the anthozoan polyp body plan is ancestral to all cnidarians (35), then the origin of strobilation (medusa formation) and its associated tissue remodeling in the stem medusozoan may have necessitated the sacrifice of the internalized tissue layers of the ancestral pharynx and mesenteries. In this case, the stem medusozoan may have overcome this loss by shifting the development of their gland cell population to the endoderm without sacrificing the selective advantage of secreting their products into the gastrovascular cavity. In support of this hypothesis, gland cells in *Hydra* are known to undergo differentiation in a location-specific manner, suggesting the identity of this cell lineage is highly sensitive to positional cues from other cells in their environment (36). Furthermore, a recent study of single-cell dynamics in *Hydra* demonstrated that gland cells acquire their identity in the endoderm only after their precursor migrates out of the ectoderm and across the mesoglea (37). Both of these studies point to the highly plastic nature of gland cell identity in *Hydra* but similar analyses in more medusozoans are needed to understand the relationship between gland cell development and cnidarian life history evolution.

## Conclusions

The transition from unicellular to multicellular life was marked by many transitions that enabled functional specialization. Unicellular taxa used trypsins for intracellular protein regulation but the origin of the regulated secretion system created new opportunities for protease activity in multiple tissue compartments. Secretion of molecules to the extracellular space enabled the development of the nervous, endocrine, immune, and digestive systems, and permitted spatial and temporal separation of multiple functions performed by a single cell. The diversification of animals was associated with a large expansion of trypsins. Trypsins with transmembrane domains first appear in the choanoflagellates but trypsins with signal peptides did not appear until the origin of animals. Subsequent duplication and divergence (e.g., through exon shuffling and retrotransposition) of genes encoding secreted proteases enabled nuanced variation in the function of these secretory cells before the increase in anatomical diversity (Figure 9).

**Figure 9.**
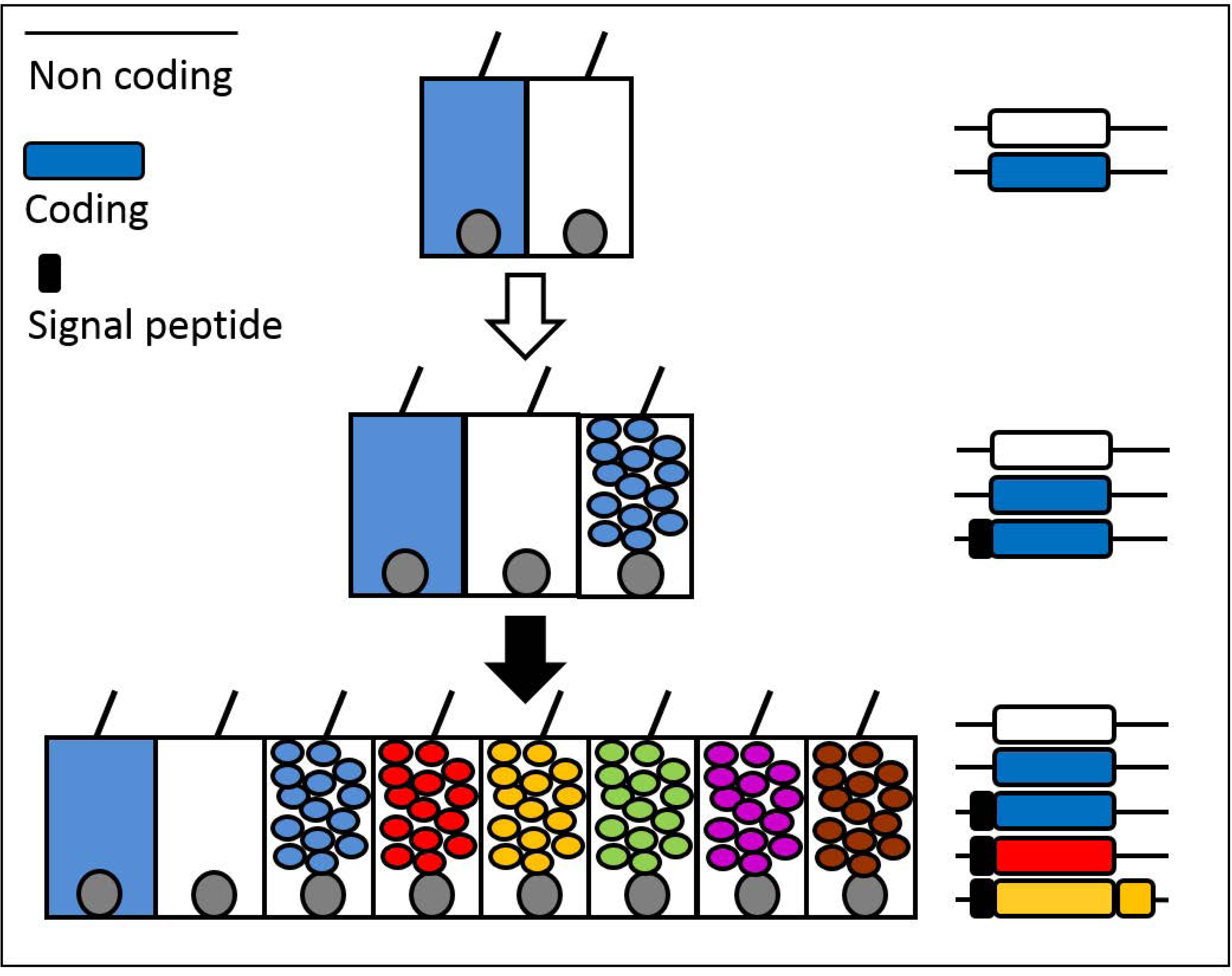
Secretory vesicles permit functional expansion without anatomical variation. Blue and white cells reflect the intracellular expression of blue and white gene products. The origin of the signal peptide directing proteins to the regulated secretion system/secretory vesicle (white arrow) permitted segregation of gene products into two distinct compartments: intracellular and intravesicular. The subsequent duplication of only few genes (black arrow) could result in the acquisition of numerous new cell types through unique and combinatorial expression. Green – co-expression of blue and yellow, purple – co-expression of blue and red, brown – co-expression of blue, yellow, and red.

## Methods

### Electron microscopy, cell proliferation assay, and in situ hybridization

Adult polyps were immobilized for 10 mins in 7.5% MgCl_2_ and processed for transmission electron microscopy as described previously (38). Samples were imaged on a Hitachi HT7700 at the University of Hawaii’s Biological Electron Microscopy facility. To identify proliferating nuclei, live polyps were incubated in 100uM EdU (in 1/3X seawater) for 30 mins at room temperature. Animals were then immobilized and fixed briefly (1.5 min) at 25C in 4% paraformaldehyde with 0.2% glutaraldehyde in phosphate buffered saline with 0.1% Tween-20 (PTw) followed by a long fixation (60 min) in 4% paraformaldehyde in PTw at 4C. Fixed tissues were analyzed using the Click-IT EdU kit (#C10340, Invitrogen, USA) following the manufacturer’s protocol. Nuclei were counter stained in a 30 min incubation in DAPI at room temperature and samples were imaged on a Zeiss 710 confocal microscope at the Whitney Lab for Marine Bioscience. To characterize the localization of target genes, we performed *in situ* hybridization following a standard protocol for *N. vectensis* (39).

### Protein domain analysis

To identify trypsin-domain proteins from *N. vectensis* we first searched the JGI protein models using the default settings with hmmsearch (HMMER 3.1b2; http://hmmer.org/) and two target HMMs: Trypsin (PF00089) and Trypsin_2 (PF13365). This approach yielded 99 putative trypsin-domain containing proteins (hereafter referred to as trypsins) with an E-value ≤ 1e-05 (40). Where multiple partial non-overlapping trypsin domains were identified from the same protein, we assumed these represented one single contiguous domain (41). Based on a reciprocal BLAST comparison with transcriptome data available publicly (11), we found 68/99 of the JGI gene models coding for trypsin proteins were incomplete. We manually corrected these sequences using the transcriptome data and used these corrected sequences for downstream analyses. We then used the transcriptome data to search protein models for evidence of pseudogenes (with premature stop codons) using the translation and alignment features in Geneious v 7.1.8 (https://www.geneious.com) and manually examined models for duplicate predictions using the JGI genome viewer. Based on these analyses, we removed 27 sequences, resulting in a final set of 72 curated trypsin protein models (FASTA file available at: https://github.com/josephryan/2019-Babonis_et_al_trypsins).

We examined the domain architecture of trypsin proteins from *N. vectensis* by searching for non-Trypsin domains in the amino acid sequences using hmmscan (HMMER 3.1b2) and the complete Pfam-A database (downloaded Oct 27, 2017). Hmmscan identifies regions of similarity between protein queries and domain models (protein profiles) derived from numerous proteins within the family from a range of animals (Bateman et al 2004). Following the protocol of Koch et al (40), we ran hmmscan using the default parameters and report only those domains with an independent (domain-specific) E-value ≤ 0.05 that were found in a protein containing a significant Trypsin (or Trypsin_2) domain. Domains that overlapped by ≤ 20% were both retained; when the overlap was >20% the domain with the lower E-value was retained. In addition to domain analysis, we manually searched an alignment of the corrected set of trypsin protein models from *N. vectensis* for the conserved residues that comprise the trypsin catalytic triad (necessary for inferring protease activity): H-57, D-102, or S-195. Finally, we searched the corrected amino acid sequences for signal peptides and transmembrane domains using SignalP v4.1 (42) and TMHMM v2.0 (43), respectively.

To characterize the origin of trypsin domain architecture, we hmmscan with the same approach described above to identify and characterize trypsins from representatives across all domains of life. We sampled three bilaterians (*Capitella teleta*, *Branchiostoma floridae*, *Homo sapiens*), ten cnidarians (*N. vectensis*, *Edwardsiella lineata*, *Aiptasia pallida*, *Anthopleura elegantissima*, *Acropora digitifera*, *Renilla renilla, Hydra magnipapillata, Calvadosia cruxmelitensis, Atolla vanhoeffeni, Alatina alata*), three non-planulozoan animals (*Mnemiopsis leidyi*, *Amphimedon queenslandica*, *Trichoplax adhaerens)*, five non-metazoan eukaryotes (*Dictyostelium discoidum, Schizosaccharomyces pombe, Capsaspora owczarzaki, Monosiga brevicolis, Salpingoeca rosetta*) and a combined database of representative archeaea and bacteria (*Candidatus aquiluna, Candidatus nitrosopumilus, Candidatus pelagibacter, Glaciecola pallidula*, *Marinobacter adhaerens,* a marine gamma proteobacterium, and a marine group I thaumarchaeote). Protein models were predicted from transcriptome data previously for *N. vectensis, E. lineata, A. pallida, A. elegantissima, A. alatina, A. vanhoeffeni*, and *P. carnea (11)*. Proteomes for *R. renilla* and *C. cruxmelitensis* were predicted from the transcriptome data reported by Kayal et al., (35) using the same methods. For all other taxa, protein models were downloaded directly (commands available at: https://github.com/josephryan/2019-Babonis_et_al_trypsins).

### Phylotocol (phylogenetic transparency)

All phylogenetic investigations were planned prior to running any analyses and all are reported in this manuscript. In most cases, these analyses were outlined beforehand in a phylotocol (44) that is posted on our GitHub site: https://github.com/josephryan/2019-Babonis_et_al_trypsins. Any analyses performed prior to being added to our phylotocol were later added to the document and justified.

### Phylogenetics

To understand the diversification of animal trypsins, we built a phylogeny using predicted proteins from *M. leidyi, A. queenslandica, T. adhaerens, N. vectensis, E. lineata, H. magnipapillata, C. teleta, B. floridae,* and *H. sapiens*. First, we used a custom script to generate alignments from these protein files using the Trypsin HMM (commands available at: https://github.com/josephryan/2019-Babonis_et_al_trypsins). All trees were constructed using a maximum likelihood framework with RAxML and IQ-TREE (45–47). We used the model finder function with IQ-TREE (-m MF) to determine the best substitution model for the alignment and then ran three approaches in parallel: RAxML with 25 parsimony starting trees, RAxML with 25 random starting trees, a single run with IQ-TREE (which, by default, uses a broad sampling of initial starting trees). We selected the best tree by comparing the maximum likelihood scores of all three approaches.

Using the Trypsin HMM we recovered 97% (70/72) of the curated trypsin proteins from *N. vectensis*. The two remaining trypsin proteins (NVJ_23745 and NVJ_203589) were recovered using the Trypsin_2 HMM. (Note: the Trypsin_2 HMM recovered only 89% (64/72) of the curated trypsins.) To understand the evolutionary relationships of these two proteins to the rest of the trypsin family, we generated another phylogeny using the same procedure as above and an alignment built using the Trypsin_2 HMM. This best tree recovered using the Trypsin_2 HMM is provided in Supplemental file 2. After inspecting both trees, we removed sequences from *B. floridae* for ease of viewing and re-ran the full analyses. All tree files and alignment files are available on our Github site (https://github.com/josephryan/2019-Babonis_et_al_trypsins).

To evaluate whether *N. vectensis* has undergone lineage-specific expansion of trypsins or if the common ancestor of all cnidarians had an equally diverse trypsin protein repertoire, we built a phylogeny of trypsin proteins from cnidarians only using a subset of the proteomes listed above. Specifically, we used four species of anthozoans (*N. vectensis, E. lineata, R. renilla, A. digitifera*) and four medusozoans (*H. magnipapillata, C. cruxmelitensis, A. vanhoeffeni, A. alata*). We then pruned all non-*Nematostella* taxa from this tree using Phyutility v.2.2.6 (48) to generate a tree for *N. vectensis* trypsins only. To examine the evolutionary history of ShK domains from *N. vectensis*, we used hmmsearch with the ShK HMM and a custom script (as above) to identify and align all ShK domains from the predicted proteome. We then used the approach described above to produce a phylogeny of ShK domains.

## Acknowledgements

We are grateful to Tina Carvahlo and Dr. Marilyn Dunlap of the Biological Electron Microscopy Facility at the University of Hawai’i Manoa for their assistance with electron microscopy. This work was supported by the National Aeronautics and Space Administration (grant NNX14AG70G to MQM) and the National Science Foundation (grant 1542597 to JFR).

## Authors’ contributions

Study design/concept: LSB, MQM, JFR; animal/tissue methods: LSB, CE; phylogenetics: LSB, JFR; other analyses: LSB; writing: LSB; review and editing: MQM, JFR, CE. All authors read and approved the final manuscript.

## Competing interests

The authors have no competing interests.

## Notes

#### Summary of Updates

New title

https://github.com/josephryan/2019-Babonis_et_al_trypsins

