## Supplementary material for "Genomic analysis of the tryptome reveals molecular mechanisms of gland cell evolution": Supplemental_file_6.pdf

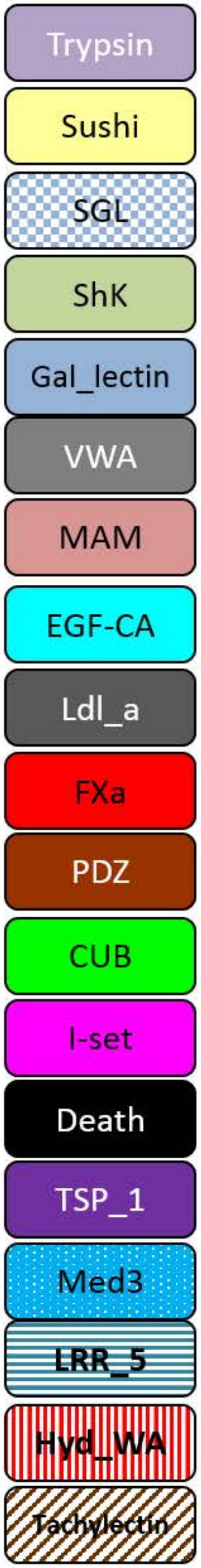


| N | Aala PID (N = 81) |
|---|---|
| 44 | See supplemental file 7 |
| 1 | m.31133 |
| 2 | m.44549, m.44547 |
| 1 | m.24221 |
| 5 | m.21496, m.51328, m.9990, m.11102, m.21497 |
| 4 | m.5584, m.3123, m.20157, m.20155 |
| 1 | m.23694 |
| 1 | m.21774 |
| 3 | m.28072, m.28077, m.28066 |
| 2 | m.52363, m.18447 |
| 5 | m.33522, m.16331, m.7126, m.33526, m.22461 |
| 1 | m.20173 |
| 3 | m.11106,, m.7124, m.11107 |
| 1 | m.8965 |
| 4 | m.39048, m.39053, m.39056, m.39050 |
| 2 | m.12633, m.12632 |
| 1 | m.4721 |

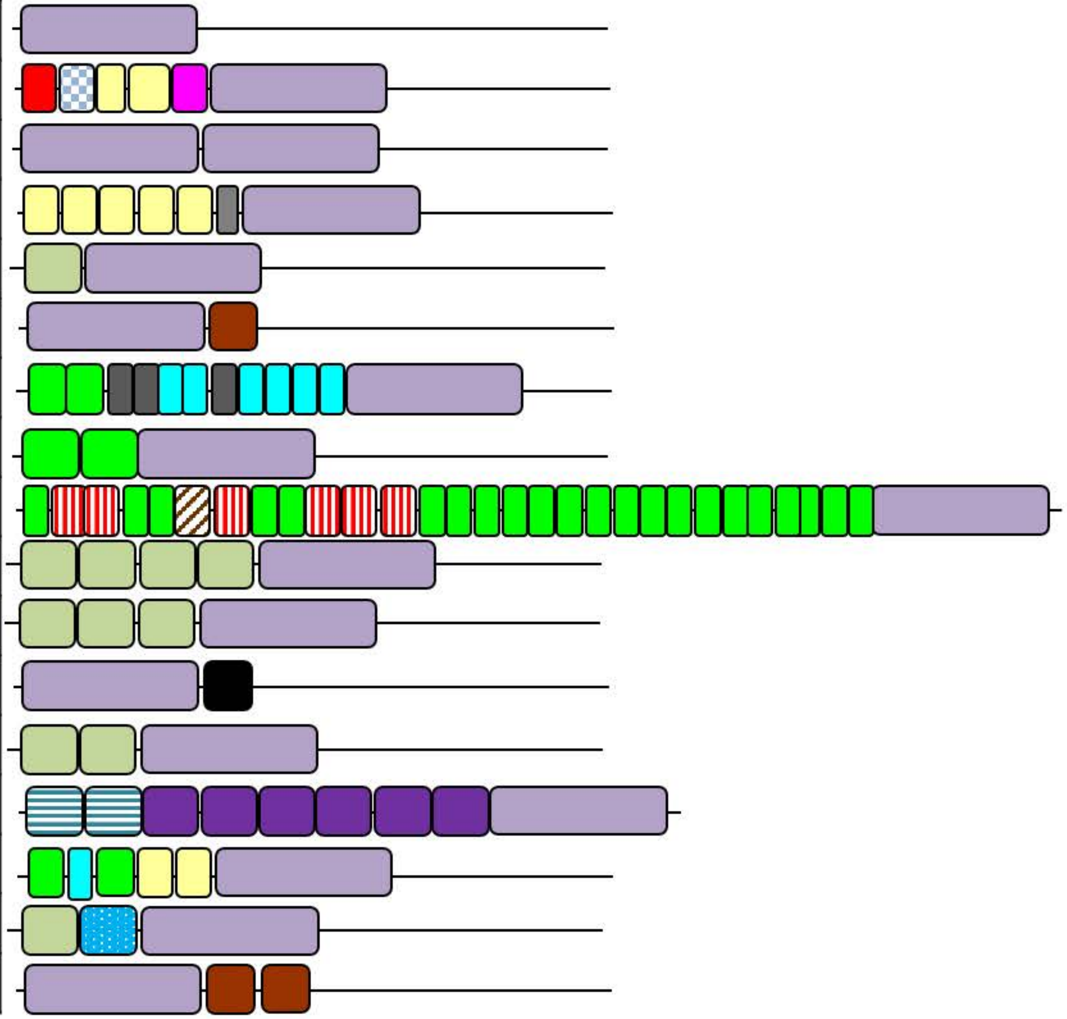

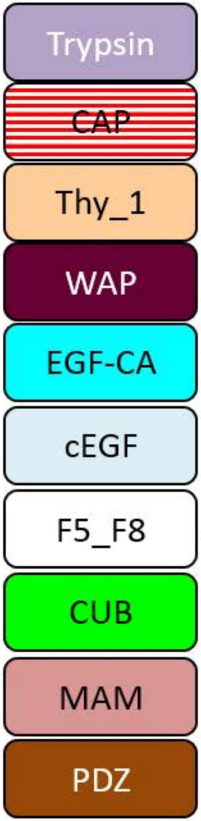


| N | Adig PID (N = 36) |
|---|---|
| **27** | **See supplemental file 7** |
| 1 | a00010306.t1 |
| 1 | a00151501.t1 |
| 1 | a00063302.t1 |
| 1 | a00161801.t1 |
| 1 | a00290801.t1 |
| 1 | a00905001.t1 |
| 2 | a01147401.t1, a00141802.t1 |
| 1 | a01851001.t1 |

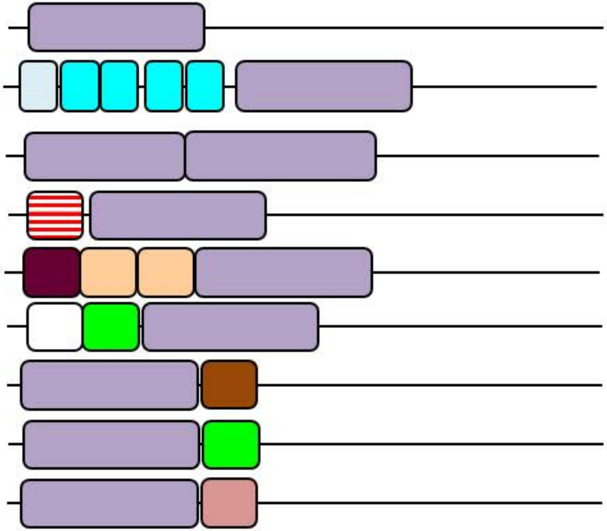

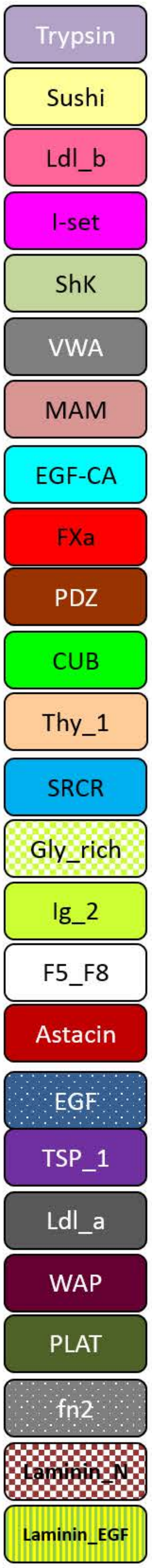


| N | Aele PID (N = 323) |
|---|---|
| **164** | **See supplemental file 7** |
| 6 | Aele.84310, Aele.84342, Aele.84327, Aele.84306, Aele.84299, Aele.84314 |
| 10 | Aele.2701, Aele.2698, Aele.2728, Aele.2699, Aele.2714, Aele.2719, Aele.2716, Aele.2697, Aele.9372, Aele.2715 |
| 3 | Aele.6394, Aele.6395, Aele.6396 |
| 3 | Aele.66398, Aele.66399, Aele.66354 |
| 31 | Aele.38170, Aele.1887, Aele.1895, Aele.82646, Aele.82645, Aele.38611, Aele.11576, Aele.38610, Aele.38171, Aele.57842, Aele.38608, Aele.73956, Aele.38609, Aele.1920, Aele.14730, Aele.1894, Aele.74104, Aele.1916, Aele.82644, Aele.57846, Aele.900, Aele.38612, Aele.11014, Aele.1902, Aele.1909, Aele.1910, Aele.1901, Aele.48490, Aele.82643, Aele.73953, Aele.1886 |
| 8 | Aele.83941, Aele.83938, Aele.83944, Aele.83942, Aele.83939, Aele.83940, Aele.83943, Aele.83945 |
| 6 | Aele.66404, Aele.66374, Aele.66379, Aele.66368, Aele.66367, Aele.66387 |
| 15 | Aele.5475, Aele.66265, Aele.14534, Aele.94292, Aele.5464, Aele.5472, Aele.14536, Aele.5451, Aele.14535, Aele.66264, Aele.5454, Aele.34635, Aele.58000, Aele.94295, Aele.66266 |
| 6 | Aele.66384, Aele.66407, Aele.66383, Aele.66373, Aele.66393, Aele.66378 |
| 10 | Aele.84388, Aele.84373, Aele.84318, Aele.84378, Aele.84383, Aele.84331, Aele.84343, Aele.84319, Aele.84355, Aele.84335 |
| 1 | Aele.6393 |
| 1 | Aele.92509 |
| 4 | Aele.97700, Aele.97702, Aele.97701, Aele.97703 |
| 2 | Aele.2727, Aele.2726 |
| 2 | Aele.66388, Aele.66394 |
| 1 | Aele.899 |
| 1 | Aele.89264 |
| 2 | Aele.81266, Aele.81267 |
| 2 | Aele.66258, Aele.66259 |
| 2 | Aele.53314, Aele.53315 |
| 2 | Aele.2720, Aele.2723 |
| 1 | Aele.25222 |
| 1 | Aele.77727 |
| 3 | Aele.48487, Aele.48486, Aele.48488 |
| 3 | Aele.17373, Aele.17374, Aele.17377 |
| 2 | Aele.62285, Aele.62282 |
| 1 | Aele.45455 |
| 1 | Aele.61797 |
| 1 | Aele.6389 |
| 2 | Aele.82630, Aele.82631 |
| 2 | Aele.89263, Aele.89262 |
| 2 | Aele.76119, Aele.76122 |
| 1 | Aele.13824 |
| 1 | Aele.56925 |
| 1 | Aele.89888 |
| 1 | Aele.89890 |
| 11 | Aele.39884, Aele.39813, Aele.39820, Aele.39774, Aele.39897, Aele.39727, Aele.39871, Aele.39821, Aele.39864, Aele.39850, Aele.39922 |
| 1 | Aele.25224 |
| 1 | Aele.34630 |
| 3 | Aele.76121, Aele.76118, Aele.76120 |
| 1 | Aele.6392 |
| 1 | Aele.92111 |

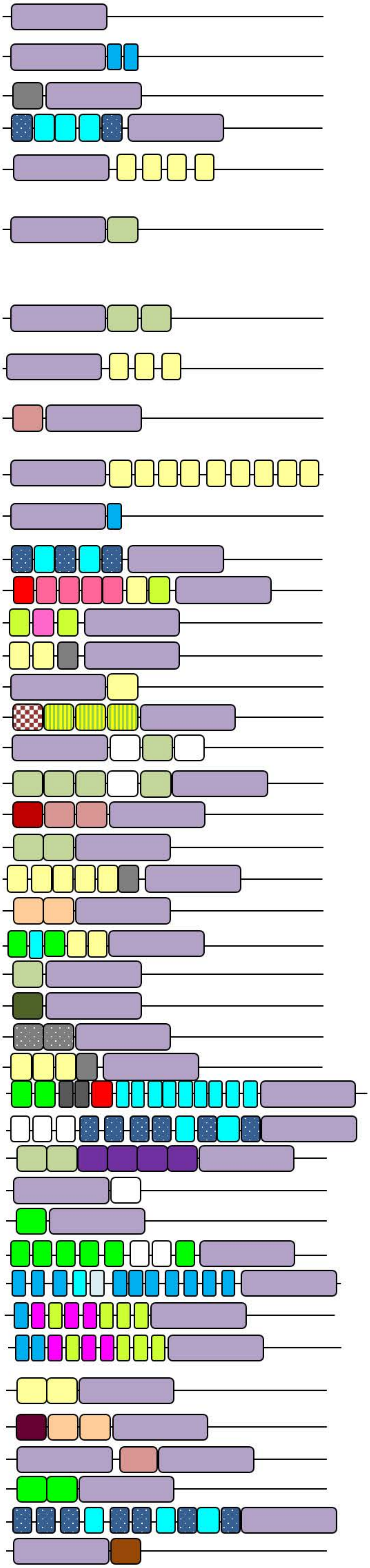

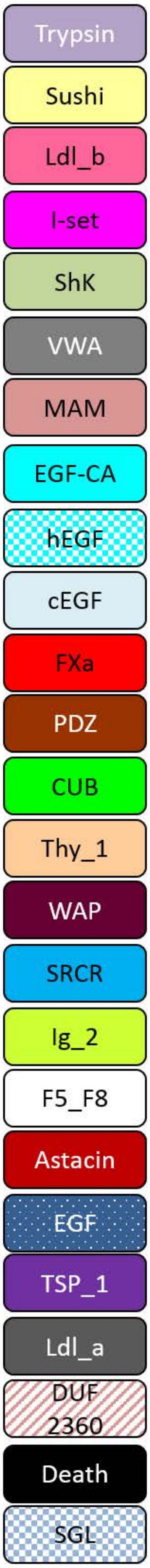


| N | Apal PID (N = 184) |
|---|---|
| 104 | See supplemental file 7 |
| 2 | m.33462, m.23664 |
| 1 | m.41150 |
| 1 | m.23666 |
| 3 | m.51059 , m.45787, m.45783 |
| 3 | m.42496, m.47728, m.42495 |
| 1 | m.36698 |
| 8 | m.6435, m.6431, m.18711, m.22591, m.825, m.6433, m.3677, m.30093 |
| 4 | m.30089, m.30095, m.989, m.988 |
| 3 | m.18982, m.18983, m.7246 |
| 1 | m.7547 |
| 18 | m.13878, m.14580, m.11058, m.21780, m.21772 ,m.12135, m.15848, m.19078, m.25132, m.21771, m.1796, m.19082, m.19076, m.21779, m.21773, m.19084, m.7481, m.6263 |
| 4 | m.21775, m.21777, m.11056, m.2682 |
| 1 | m.25742 |
| 2 | m.38156, m.38152 |
| 2 | m.41152, m.25543 |
| 1 | m.16467 |
| 2 | m.11299, m.11301 |
| 2 | m.6264, m.7993 |
| 1 | m.45967 |
| 2 | m.19137, m.19134 |
| 1 | m.35972 |
| 1 | m.26197 |
| 1 | m.35973 |
| 2 | m.25497, m.25498 |
| 1 | m.44400 |
| 1 | m.28664 |
| 1 | m.25542 |
| 1 | m.26062 |
| 1 | m.33460 |
| 1 | m.29764 |
| 1 | m.14768 |
| 1 | m.12546 |
| 2 | m.25738, m.25740 |
| 1 | m.37592 |
| 1 | m.12721 |
| 1 | m.30094 |

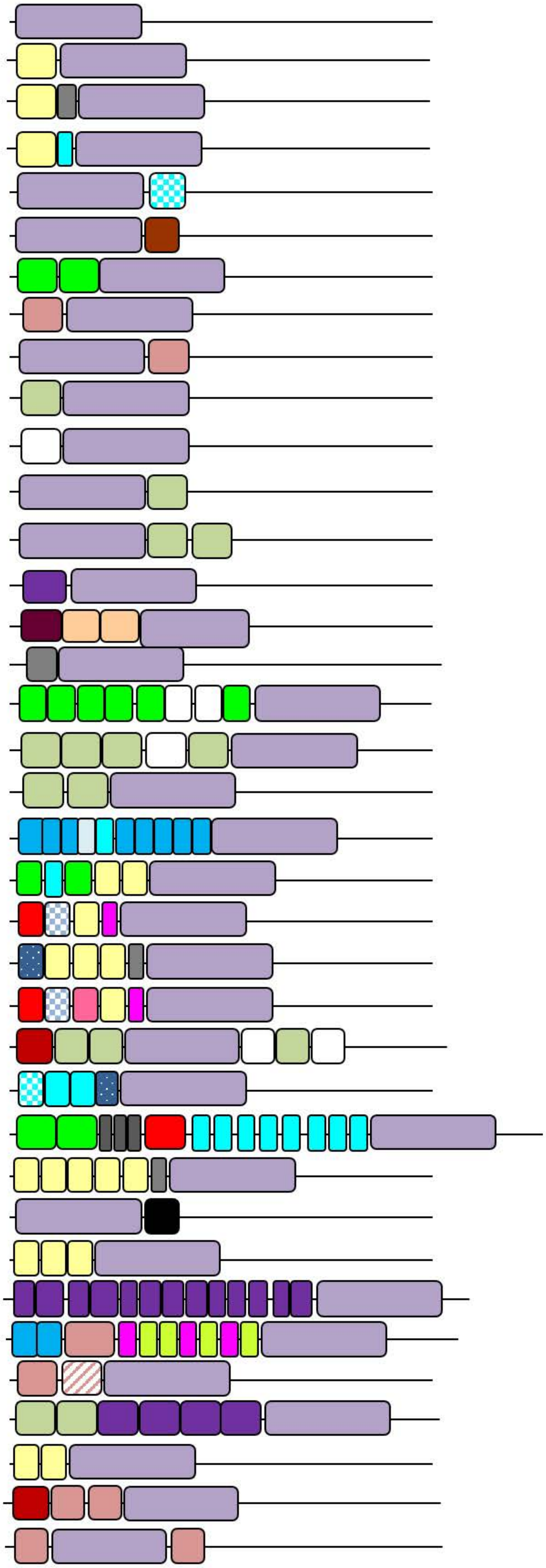

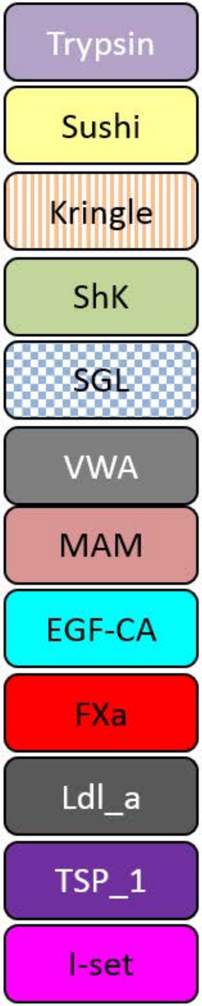


| N | Avan PID (N=48) |
|---|---|
| 26 | **See supplemental file 7** |
| 2 | **m.14473, m.14474** |
| 1 | **m.13572** |
| 4 | **m.13793, m.13799, m.13795, m.13798** |
| 2 | **m.16833, m.16835** |
| 2 | **m.16324, m.16325** |
| 2 | **m.12355, m.12356** |
| 2 | **m.14377, m.14378** |
| 2 | **m.13573, m.13571** |
| 1 | **m.13469** |
| 1 | **m.12856** |
| 2 | **m.14376, m.1092** |
| 1 | **m.6618** |

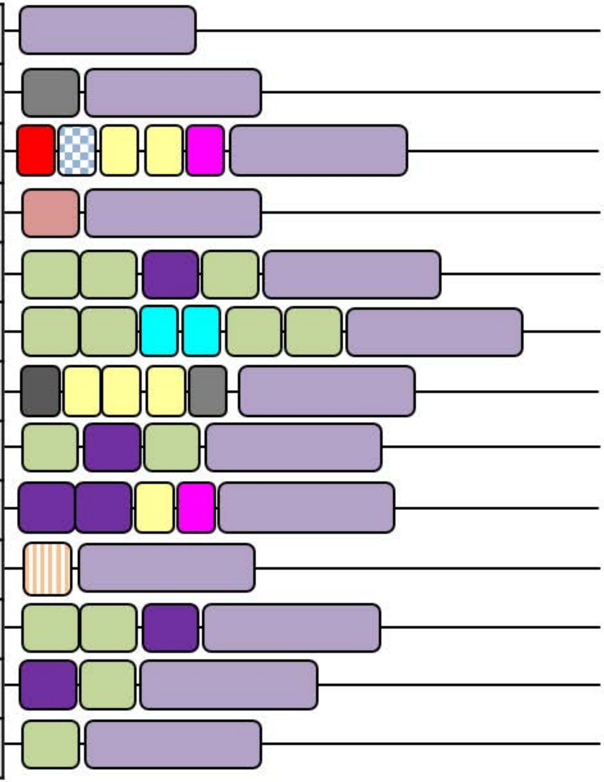

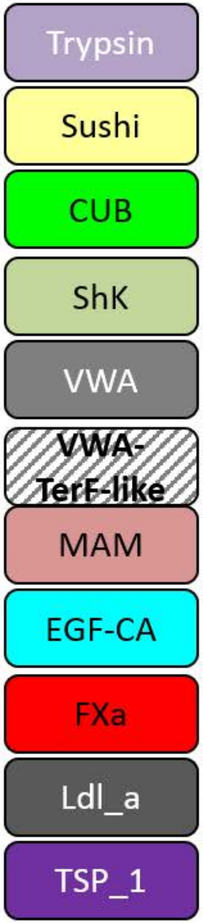


| N | Ccrux PID (N = 66) |
|---|---|
| **40** | **See supplemental file 7** |
| 2 | Ccrux_8092, Ccrux_8093 |
| 4 | Ccrux_46743, Ccrux_46742, Ccrux_46741, Ccrux_46740 |
| 3 | Ccrux_47302, Ccrux_47295, Ccrux_47298 |
| 2 | Ccrux_9402, Ccrux_29728 |
| 1 | Ccrux_62140 |
| 1 | Ccrux_47301 |
| 2 | Ccrux_1715, Ccrux_29729 |
| 3 | Ccrux_37615, Ccrux_3424, Ccrux_37613 |
| 1 | Ccrux_20375 |
| 1 | Ccrux_38926 |
| 1 | Ccrux_38929 |
| 1 | Ccrux_38934 |
| 3 | Ccrux_26746, Ccrux_64449, Ccrux_27223 |
| 1 | Ccrux_2261 |

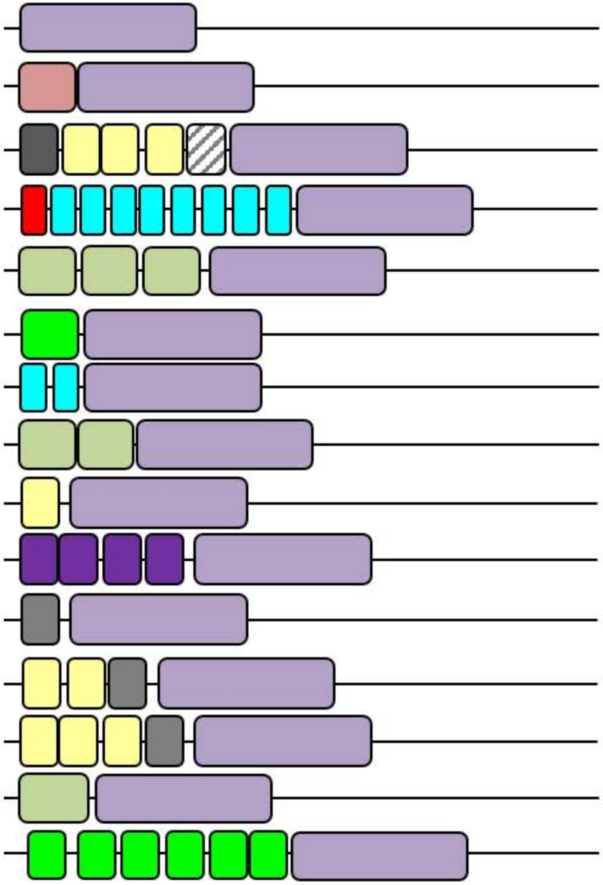

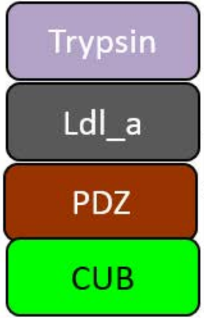


| N | Ctel PID (N=47) |
|---|---|
| **43** | **See supplemental file 7** |
| 1 | 93602 |
| 1 | 201465 |
| 1 | 228376 |
| 1 | 176051 |

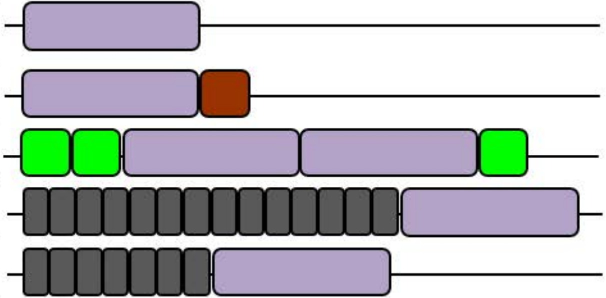

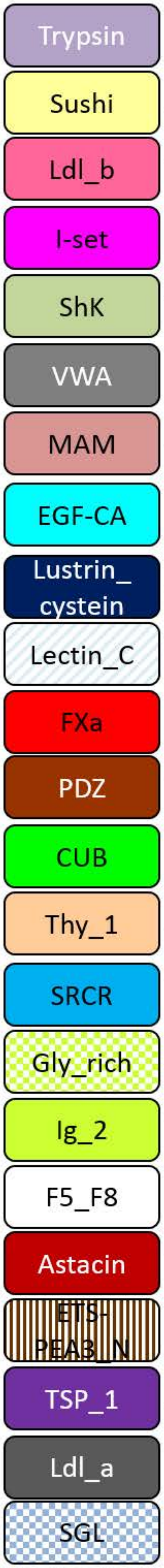


| N | Elin PID (N = 89) |
|---|---|
| **43** | **See supplemental file 7** |
| 1 | m.3862 |
| 1 | m.26131 |
| 1 | m.13004 |
| 1 | m.29106 |
| 7 | m.12619, m.9962, m.1810, m.6939, m.23171, m.17400, m.28139 |
| 1 | m.11596 |
| 1 | m.21570 |
| 2 | m.30277, m.5760 |
| 1 | m.21993 |
| 1 | m.13972 |
| 6 | m.13934, m.996, m.13941 , m.42346, m.13937, m.13938 |
| 3 | m.14356, m.14360, m.14355 |
| 2 | m.6288, m.6285 |
| 1 | m.4322 |
| 2 | m.3665, m.13971 |
| 1 | m.13939 |
| 1 | m.15907 |
| 1 | m.28621 |
| 1 | m.31483 |
| 2 | m.9787, m.9780 |
| 1 | m.6310 |
| 2 | m.10541, m.10547 |
| 1 | m.6241 |
| 1 | m.31802 |
| 1 | m.41352 |
| 1 | m.10532 |
| 1 | m.20616 |
| 1 | m.13584 |

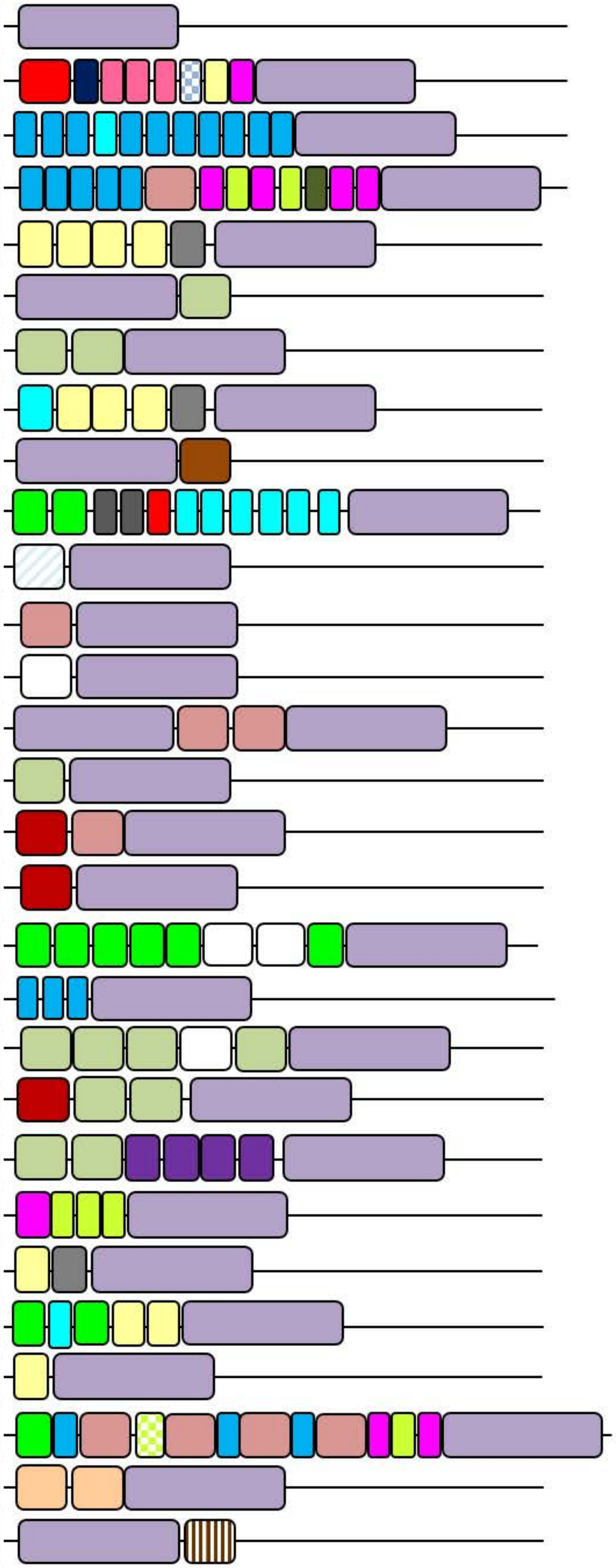

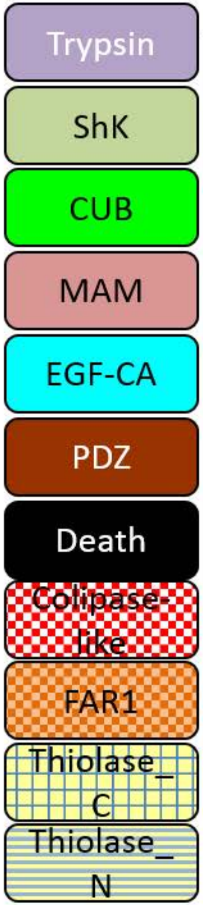


Hmag PID (N = 27)

| N | |
|---|---|
| **11** | **See supplemental file 7** |
| 1 | XP_002161936.2 |
| 1 | XP_002168460.2 |
| 1 | XP_002154014.2 |
| 7 | XP_002162561.1, XP_002168881.2, XP_002164783.2, XP_002166222.2, XP_002158991.2, XP_002163360.2, XP_002164216.1 |
| 1 | XP_002161356.1 |
| 1 | XP_004206897.1 |
| 1 | XP_002160968.2 |
| 2 | XP_002154609.2, XP_004206333.1 |
| 1 | XP_002168857.2 |

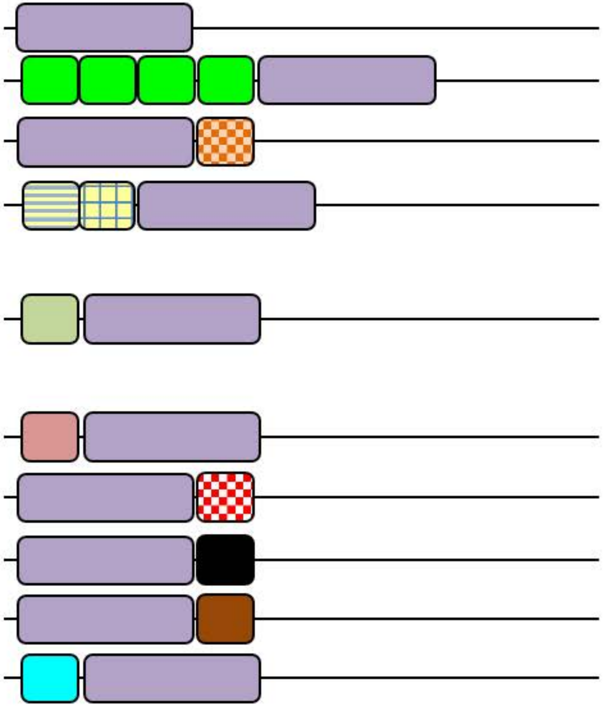

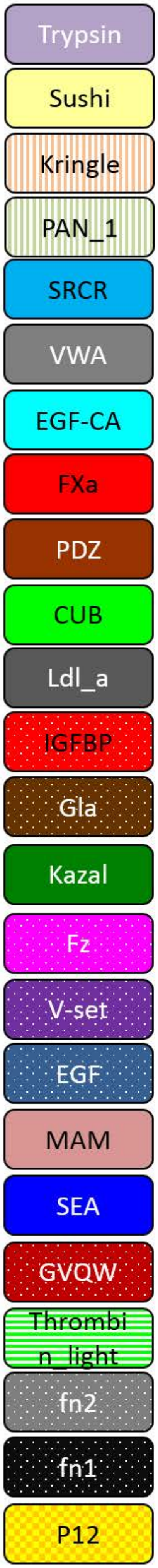


| N | Hsap PID (N = 124) |
|---|---|
| 66 | See supplemental file 7 |
| 2 | NP_000496.2, NP_001284368.1 |
| 1 | NP_001701.2 |
| 1 | XP_006713229.1 |
| 1 | NP_037379.1 |
| 1 | NP_001245219.1 |
| 1 | XP_011544121.1 |
| 1 | XP_011514713.1 |
| 5 | NP_000495.1, XP_005263772.1, NP_000122.1, NP_000124.1, XP_011535827.1 |
| 5 | NP_872308.2, NP_001107859.1, NP_054777.2, NP_004253.1, NP_997290.2 |
| 1 | NP_002649.1 |
| 2 | NP_892028.1, NP_001275681.1 |
| 1 | NP_001128571.1 |
| 4 | NP_006601.2, NP_001724.3, NP_001870.3, NP_958850.1 |
| 3 | NP_001193718.1, NP_001243246.1, XP_005271670.1 |
| 3 | NP_002766.1, NP_444272.1, NP_710159.1 |
| 1 | NP_004123.1 |
| 1 | NP_002763.2 |
| 1 | NP_003610.2 |
| 2 | NP_000292.1, NP_001258662.1 |
| 1 | NP_001269605.1 |
| 1 | NP_001171534.1 |
| 1 | NP_001010932.1 |
| 1 | NP_000497.1 |
| 1 | XP_011518939.1 |
| 1 | NP_705837.1 |
| 1 | NP_127509.1 |
| 1 | NP_005568.2 |
| 1 | NP_001265514.1 |
| 2 | XP_011530232.1, XP_005262878.1 |
| 1 | NP_068813.1 |
| 1 | XP_011511056.1 |
| 1 | XP_011526280.1 |
| 1 | XP_011518365.1 |
| 1 | XP_011530222.1 |

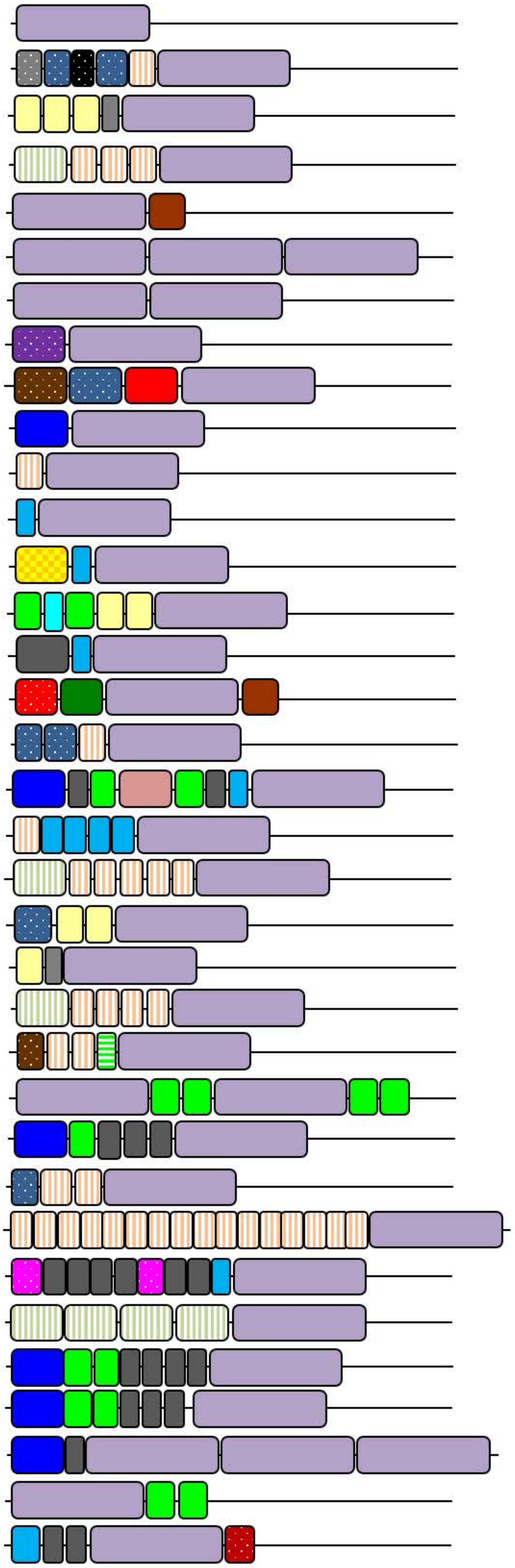

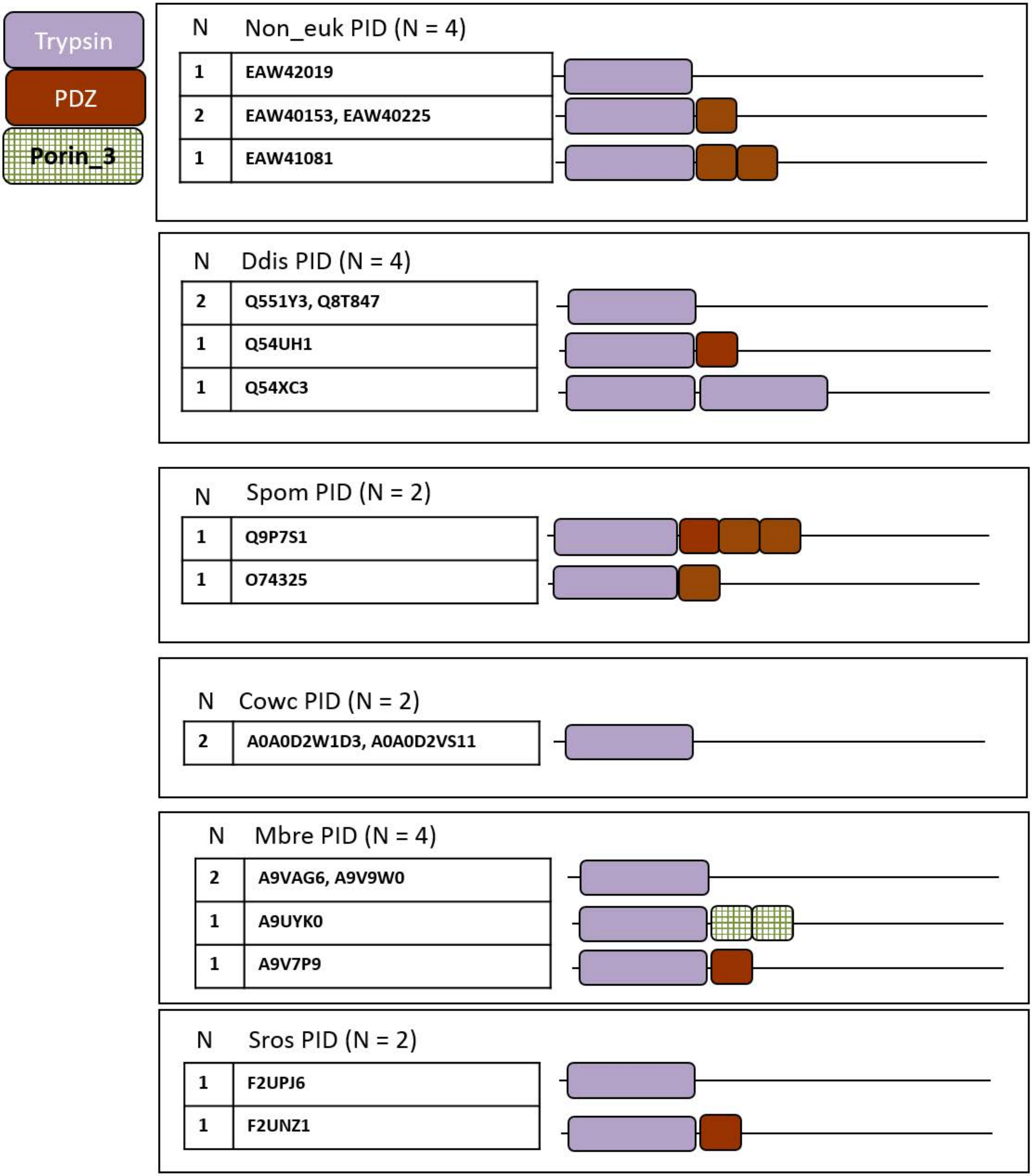

Trypsin
PDZ
Porin_3
N
Non_euk PID (N = 4)
1
EAW42019
2
EAW40153, EAW40225
1
EAW41081
N
Ddis PID (N = 4)
2
Q551Y3, Q8T847
1
Q54UH1
1
Q54XC3
N
Spom PID (N = 2)
1
Q9P7S1
1
O74325
N
Cowc PID (N = 2)
2
A0A0D2W1D3, A0A0D2VS11
N
Mbre PID (N = 4)
2
A9VAG6, A9V9W0
1
A9UYK0
1
A9V7P9
N
Sros PID (N = 2)
1
F2UPJ6
1
F2UNZ1

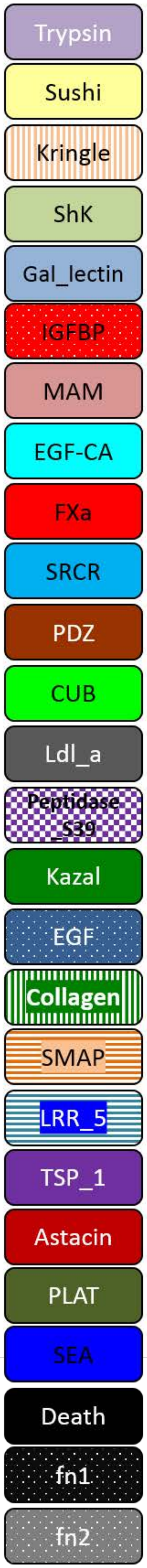


| N | Pcar PID (N = 110) |
|---|---|
| **62** | **See supplemental file 7** |
| 3 | m.43916, m.43917, m.43918 |
| 1 | m.16027 |
| 3 | m.38345, m.38348, m.3451 |
| 1 | m.58863 |
| 1 | m.28455 |
| 1 | m.8768 |
| 4 | m.37166, m.12131, m.35147, m.37168 |
| 1 | m.38794 |
| 2 | m.53227, m.53221 |
| 7 | m.29647, m.506, m.70178, m.1817, m.9828, m.12677, m.38773 |
| 1 | m.11211 |
| 2 | m.58252, m.58254 |
| 1 | m.31973 |
| 2 | m.48826, m.48828 |
| 3 | m.3516, m.3515, m.38990 |
| 2 | m.29104, m.29107 |
| 2 | m.36016, m.11427 |
| 4 | m.32836, m.55298, m.36020, m.26179 |
| 1 | m.28454 |
| 1 | m.23319 |
| 1 | m.22580 |
| 1 | m.24689 |
| 1 | m.16474 |
| 1 | m.4072 |
| 1 | m.2429 |

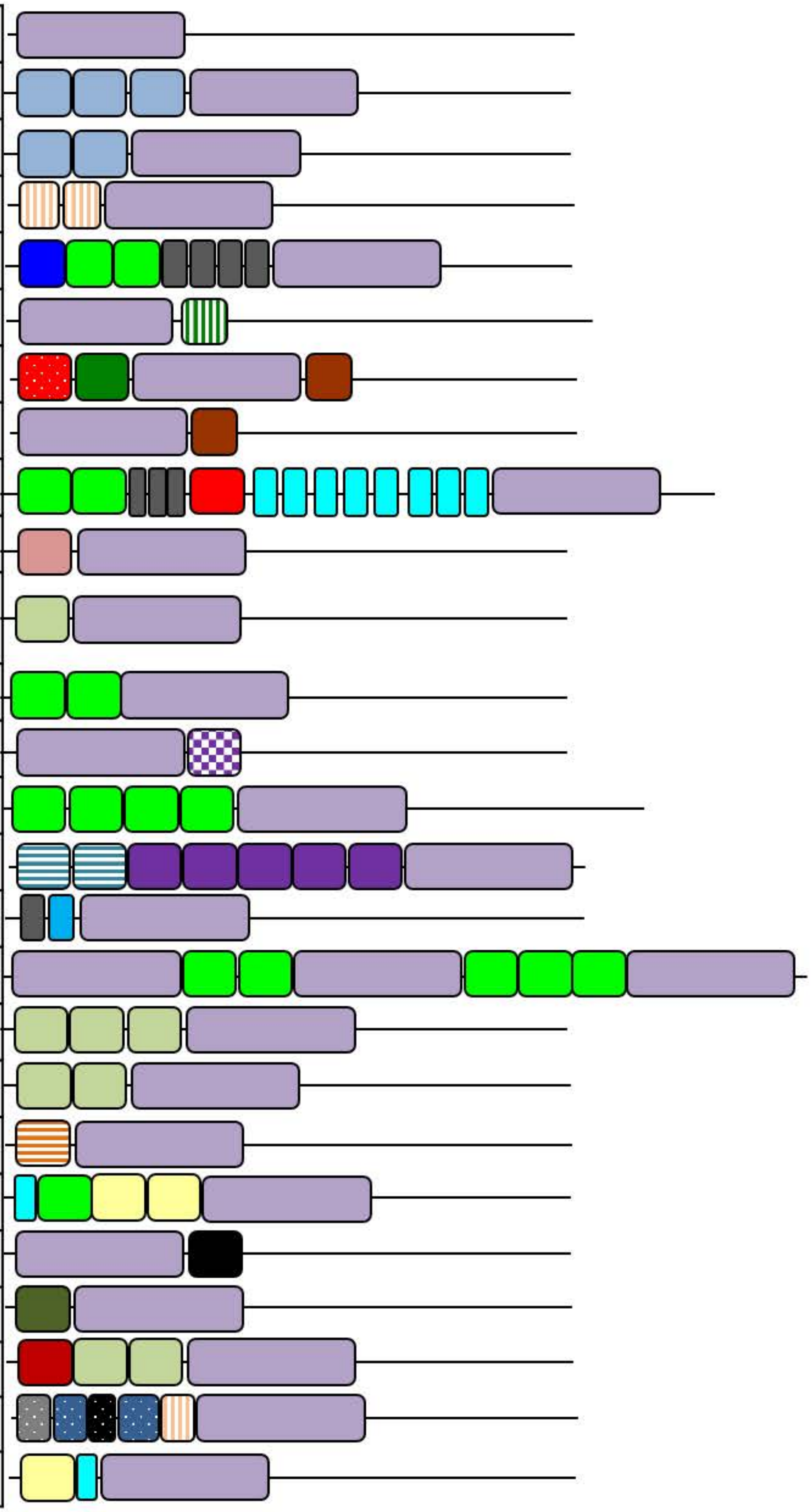

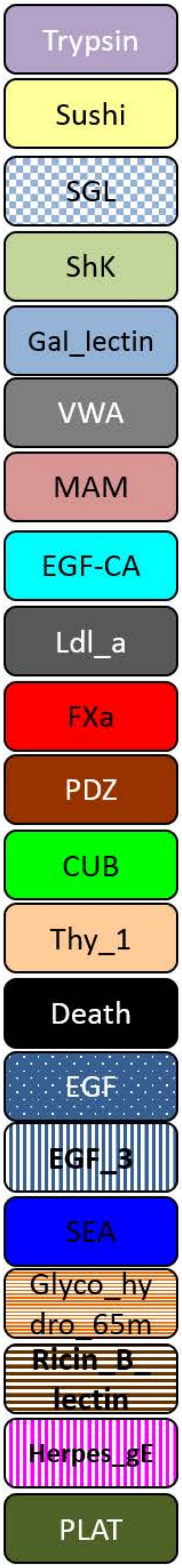


| N | Rren PID (N = 54) |
|---|---|
| **32** | **See supplemental file 7** |
| 1 | Reni_reni.v2.21181 |
| 1 | Reni_reni.v2.09705 |
| 1 | Reni_reni.v2.61890 |
| 1 | Reni_reni.v2.01988 |
| 1 | Reni_reni.v2.06917 |
| 1 | Reni_reni.v2.33190 |
| 1 | Reni_reni.v2.29630 |
| 1 | Reni_reni.v2.29657 |
| 1 | Reni_reni.v2.02803 |
| 1 | Reni_reni.v2.54597 |
| 1 | Reni_reni.v2.31223 |
| 1 | Reni_reni.v2.35127 |
| 1 | Reni_reni.v2.64201 |
| 1 | Reni_reni.v2.22398 |
| 1 | Reni_reni.v2.05955 |
| 1 | Reni_reni.v2.19590 |
| 1 | Reni_reni.v2.51451 |
| 1 | Reni_reni.v2.11546 |
| 1 | Reni_reni.v2.49481 |
| 1 | Reni_reni.v2.28079 |
| 1 | Reni_reni.v2.14471 |
| 1 | Reni_reni.v2.39164 |

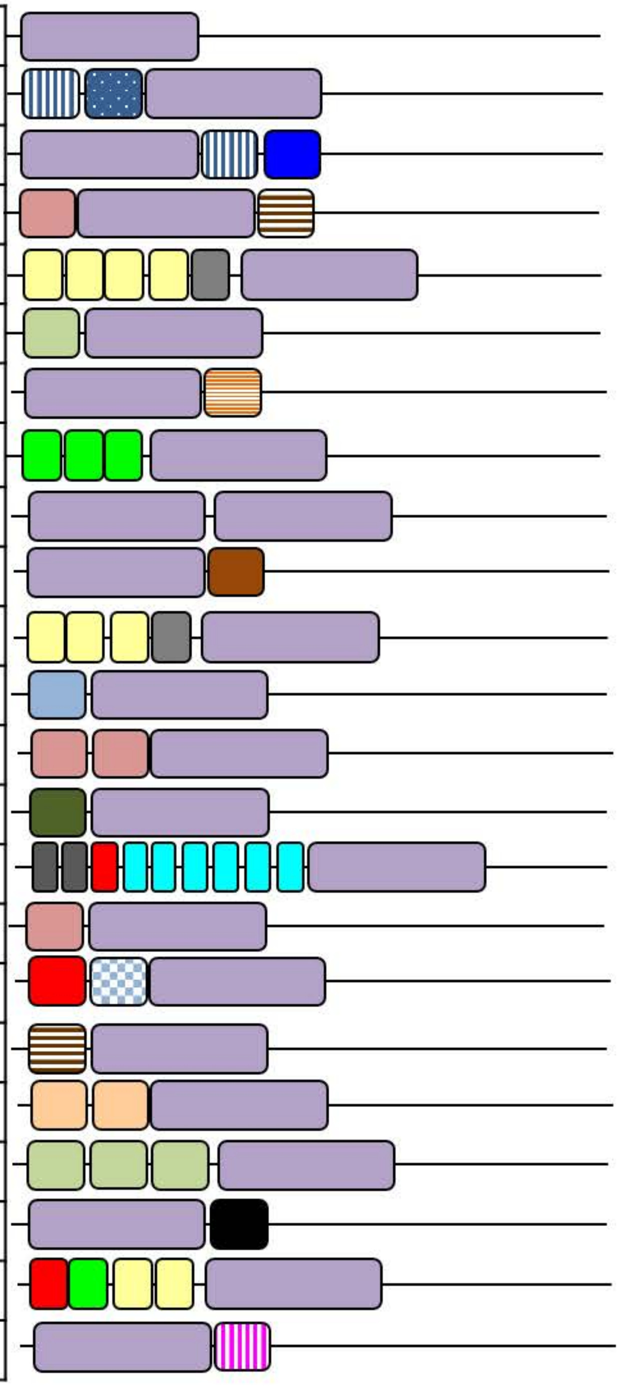

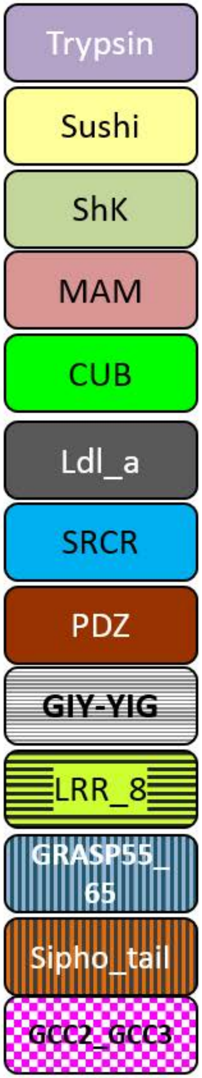


| N | Tad PID (N = 25) |
|---|---|
| **21** | **See supplemental file 7** |
| 1 | 53232 |
| 1 | 53235 |
| 1 | 55983 |
| 1 | 58565 |

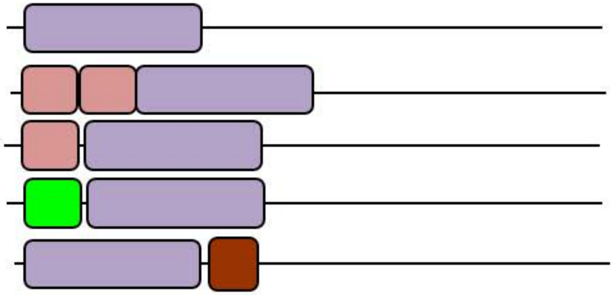

Aque PID (N = 3)

| | |
|---|---|
| 2 | XP_003387666.1, XP_003390728.1 |
| 1 | XP_003386366.1 |

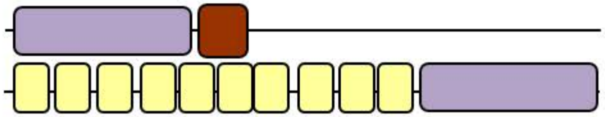

Mlei Protein ID (N=20)

| | |
|---|---|
| **6** | **See supplemental file 7** |
| 6 | ML15095a, ML033911a, ML06704a, ML006112a, ML00576a, ML002233a |
| 1 | ML11643a |
| 1 | ML216323a |
| 1 | ML142413a |
| 1 | ML279621a |
| 1 | ML223530a |
| 1 | ML398319a |
| 1 | ML00923a |
| 1 | ML00717a |

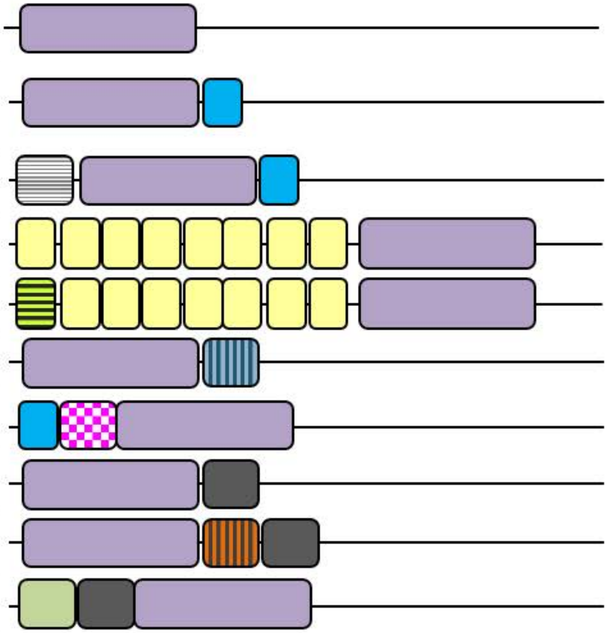
