## Supplementary figures and images for "Genomic analysis of the tryptome reveals molecular mechanisms of gland cell evolution"

### Supplemental_file_1.pdf

Supplemental file 1

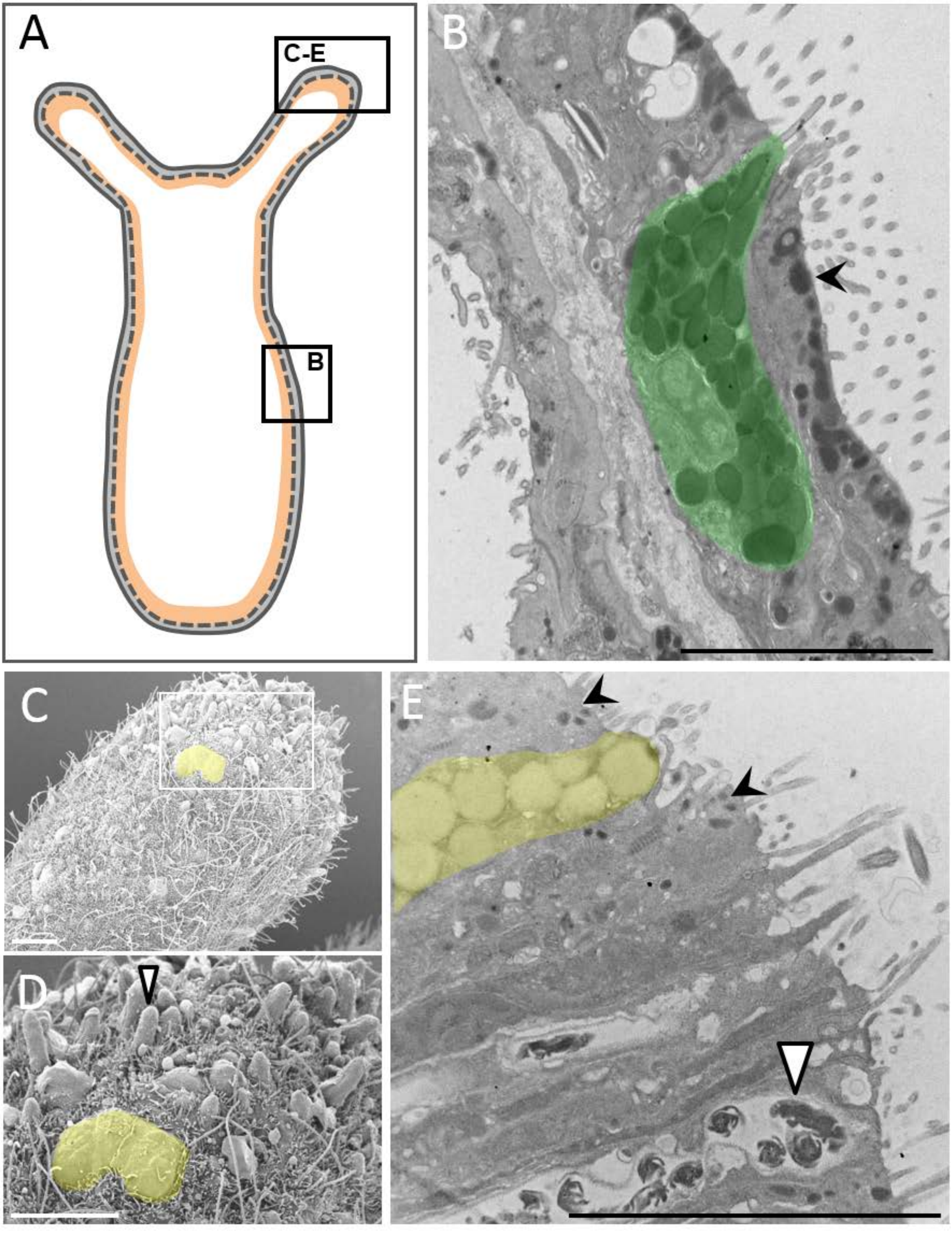

### Supplemental_file_2.pdf

Supplemental file 2

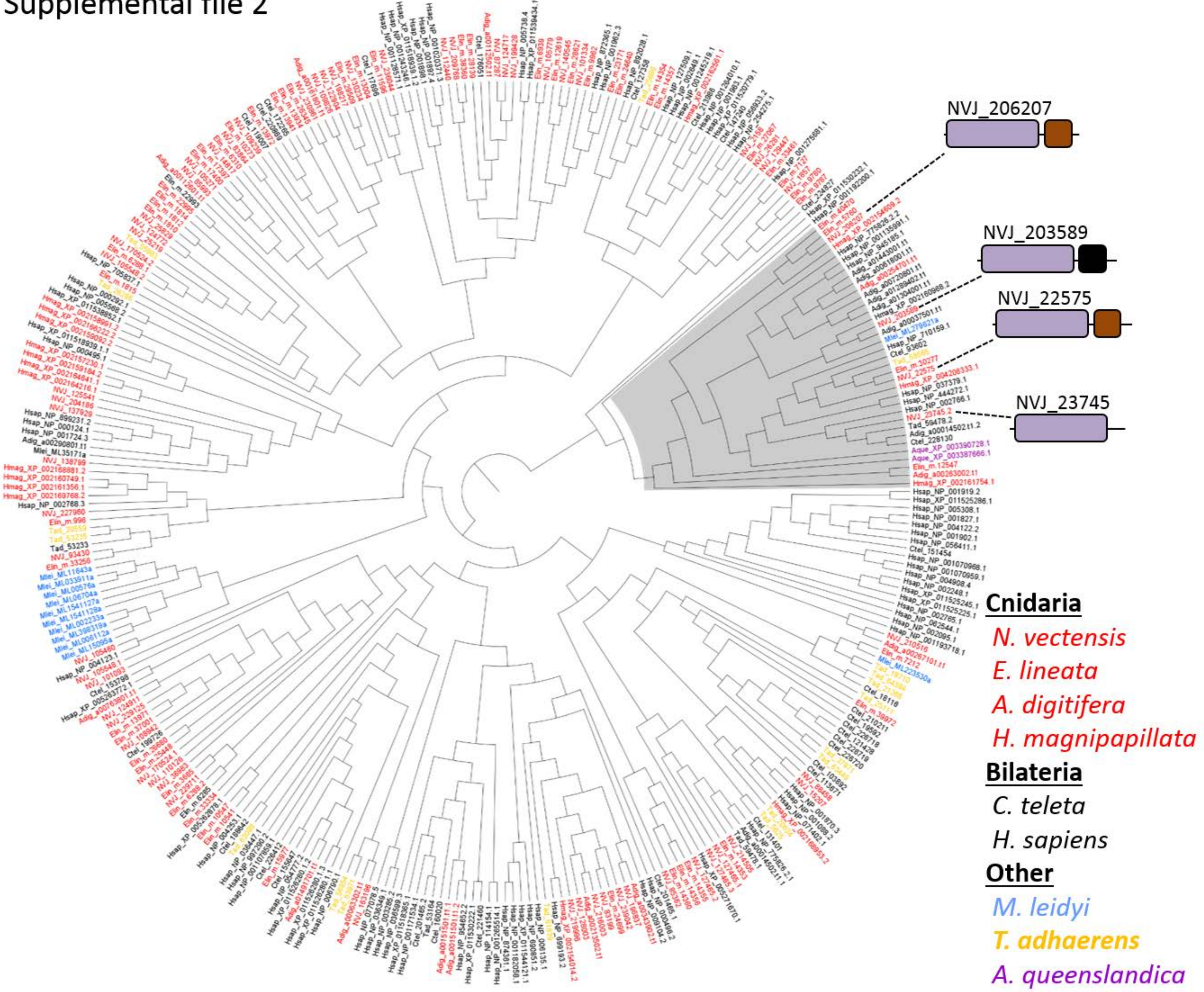

### Supplemental_file_3.pdf

Supplemental file 3

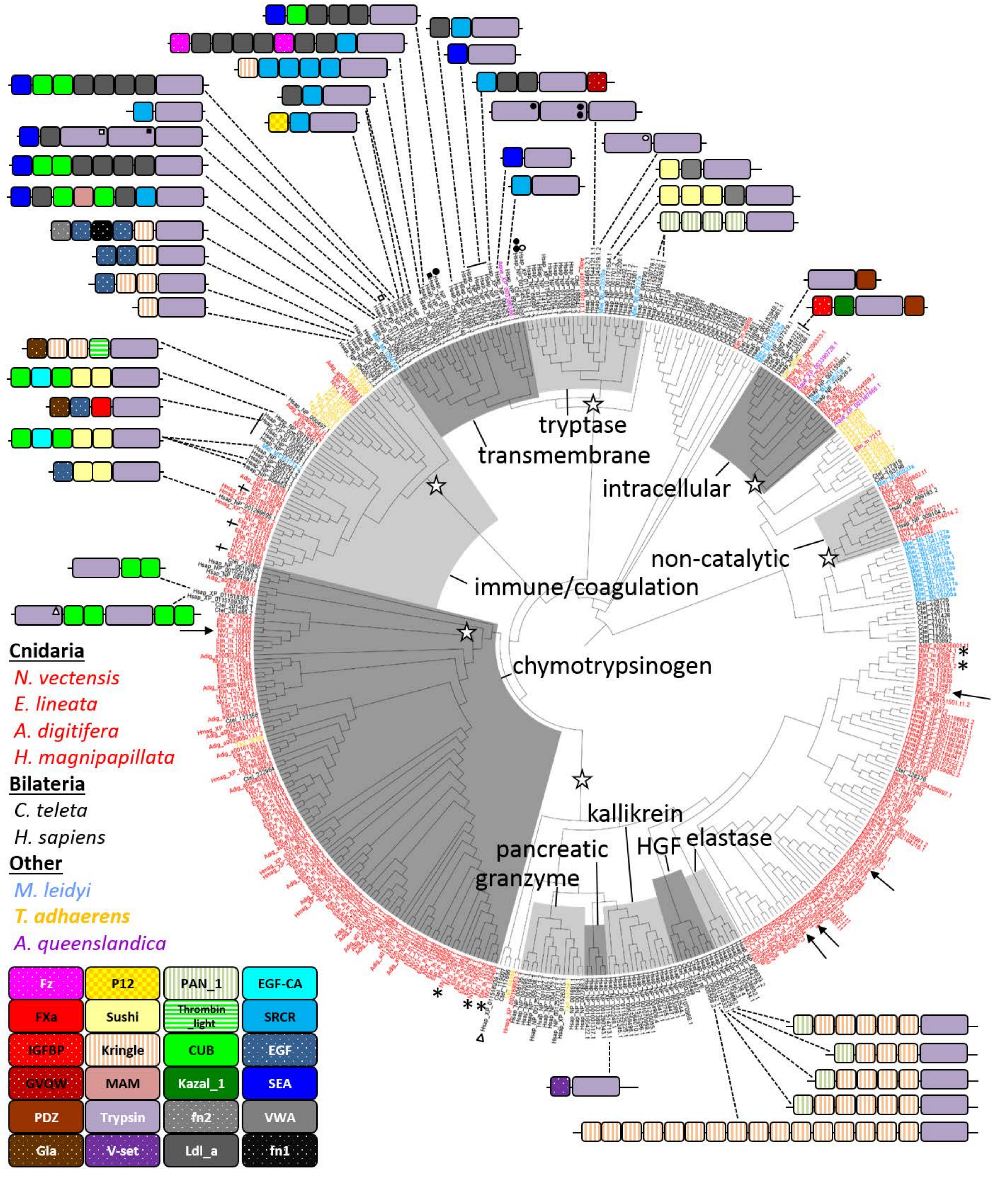

### Supplemental_file_5.pdf

Supplemental file 5

| | NVJ Protein ID |
|---|---|
| **26** | **...** |
| 2 | 128003, 216003 |
| **2** | 22575, 206207 |
| **1** | 98319, |
| **1** | 137929 |
| **1** | 41116 |
| **1** | 204186 |
| **1** | 125541 |
| **1** | 138799 |
| **1** | 109239 |
| **1** | 199744 |
| **1** | 127465 |
| **1** | 85345 |
| **4** | 93430, 229711, 110126, 227960 |
| **1** | 200868 |
| **1** | 170524 |
| **1** | 105548 |
| **2** | 218669, 218670 |
| **2** | 236044, 229125 |
| **1** | 112683 |
| **9** | 214505, 112440, 140545, 2158, 209768, 105779, 124717, 101334, 14817 |
| **1** | 25829 |
| **1** | 1857 |
| **1** | 105271 |
| **1** | 107554 |
| **1** | 85993 |
| **1** | 203589 |
| **1** | 101093 |
| **1** | 164017 |
| **1** | 109826 |
| **1** | 163196 |
| **1** | 199428 |

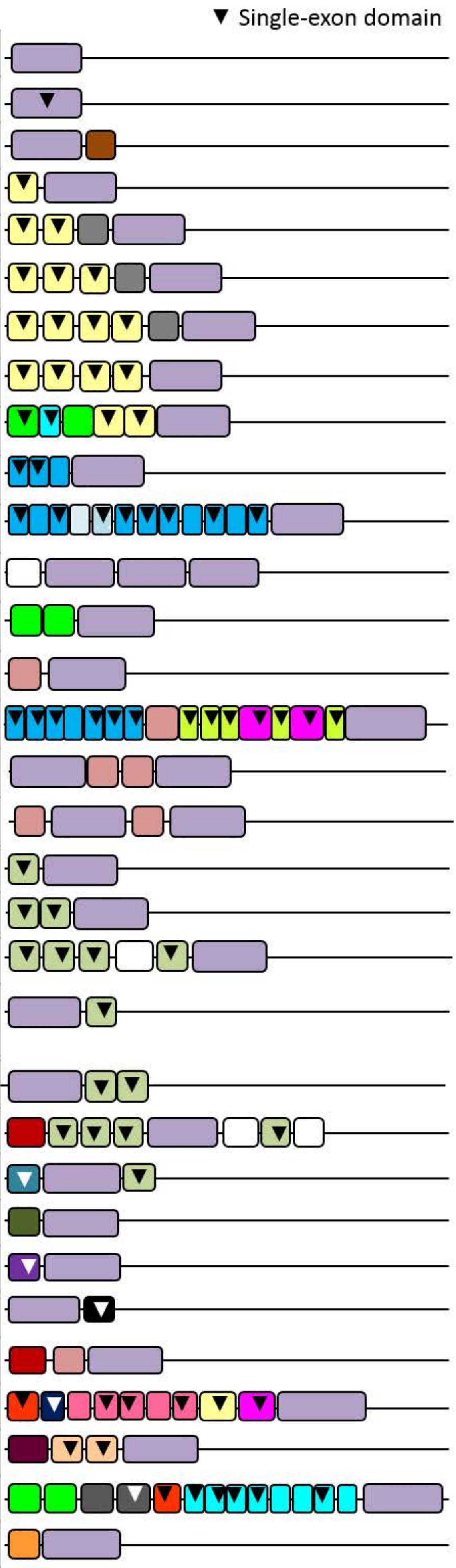
